# A claudin5-binding peptide enhances the permeability of the blood-brain-barrier

**DOI:** 10.1101/2024.04.29.591687

**Authors:** Martina Trevisani, Alessandro Berselli, Giulio Alberini, Eleonora Centonze, Silvia Vercellino, Veronica Cartocci, Enrico Millo, Dinu Zinovie Ciubanu, Andrea Armirotti, Francesco Pisani, Federico Zara, Valentina Castagnola, Luca Maragliano, Fabio Benfenati

## Abstract

The blood-brain barrier (BBB) is essential to maintain brain homeostasis and healthy conditions but it also prevents drugs from reaching brain cells. In the BBB, tight junctions (TJs) are multi-protein complexes located at the interface between adjacent brain endothelial cells that regulate paracellular diffusion and claudin-5 (CLDN5) is the major component of the TJ portfolio, playing a pivotal role in restricting the paracellular traffic. In view of obtaining fine control over the transport across the BBB, the use of competing peptides able to bind CLDN5 to induce transient and regulated permeabilization of the paracellular passage is emerging as a potentially translatable strategy for clinical applications. In this work, we designed and tested short peptides with improved solubility and biocompatibility using a combined approach that involved structural modeling techniques and *in vitro* validation, generating a robust workflow for the design, screening, and optimization of peptides for the modulation of the BBB paracellular permeability. We designed a selection of 11- to 16-mer compounds derived from the first CLDN5 extracellular domain and from the CLDN5-binding domain of *Clostridium perfringens* enterotoxin and determined their efficiency in enhancing BBB permeability. The computational analysis classified all tested peptides based on solubility and affinity to CLDN5, and provided atom-level details of the binding process. From our screening, we identified a novel CLDN5-derived peptide, here called *f1-C5C2*, which demonstrated good solubility in biological media, efficient binding to CLDN5 subunits, and capability to increase permeability at low concentrations. The peptidomimetic *in silico/in vitro* strategy described here can achieve a transient and reversible permeabilization of the BBB with potential applications in the pharmacological treatment of brain diseases.

**HIGHLIGHTS:**

- Water-soluble peptidomimetics are used to competitively bind claudin-5 tight junction proteins and increase the permeability of the blood-brain barrier;
- Trans-endothelial electrical resistance and dissociation constant measurements demonstrate the binding affinity of the peptide *f1-C5C2* for claudin-5;
- Unbinding free energy calculations correlated with experimental results and provided information on the protein-peptide binding interface.
- Incubation with the peptide *f1-C5C2* allows paracellular transport of 4K, but not 70K, dextran.

**GRAPHICAL ABSTRACT:** 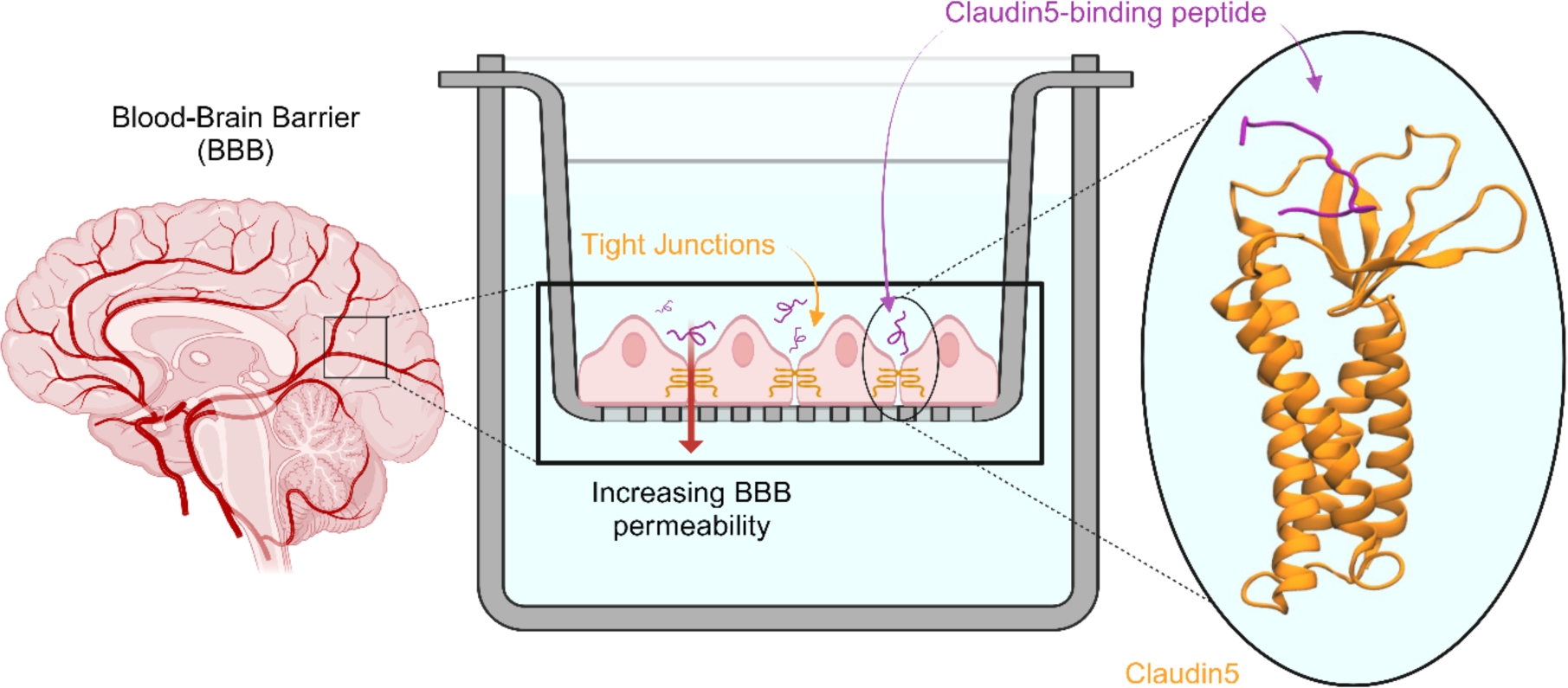

## INTRODUCTION

The blood-brain barrier (BBB) represents the primary defense layer for the brain and has a pivotal role in maintaining its homeostasis. However, it also hinders therapeutic interventions in case of brain disorders, such as neurodegeneration, glioblastoma, and various genetic diseases [1,2]. As a notable example, in the glucose transporter-1 deficiency syndrome (GLUT1DS) a loss-of-function mutation in the SLC2A1 gene encoding for the unique Glut1 transporter in brain endothelial cells prevents the appropriate supply of glucose to maintain physiological brain functions [3–5].

Because of these limitations, there is a high demand for innovative approaches to regulate BBB permeability under pathological conditions [1,6–10]. Many strategies are under investigation to increase the passage of small molecules across the BBB, by facilitating either their crossing of the endothelial cells (the transcellular route) or their passage between adjacent endothelial cells (the paracellular route) [6,11–18]. The BBB paracellular pathway is controlled by specialized protein assemblies called tight junctions (TJs), whose backbone is formed by members of the claudin (CLDN) family [19,20]. Their topology is characterized by a four-transmembrane helix bundle (TM1-4) with two extracellular loops (ECL1-2), cytoplasmic N/C termini, and an intracellular loop [19–22]. To form TJs, CLDN monomers associate in strands within the lateral membrane of endothelial cells, while strands from neighboring cells associate in *trans* across the paracellular space. CLDN5 is the most expressed member of the family in the TJs of the BBB and has a key role in restricting the paracellular traffic of molecules and ions [1,22].

Inspired by studies on CLDN1-[14,23] and CLDN4-[24,25] expressing cells, efforts have been recently devoted to studying molecular approaches based on competing peptides able to modulate the formation of CLDN5 multimers. Two families of peptides have been investigated, including (i) derivatives of the non-toxic C-terminal domain of *Clostridium perfringens* enterotoxin (cCPE) [12,26,27] and (ii) a peptidomimetic based on the murine CLDN5 (mCLDN5) ECL1 domain, known as C5C2 [11,27]. Despite these seminal endeavors, the design of CLDN-modulating peptides has been hindered by the limited structural information available on these proteins. Indeed, although the structure of other CLDNs in the monomeric form has been determined experimentally [28–31], that of CLDN5 is not yet available. Moreover, there are still major gaps in our understanding of the assembly of CLDNs in TJs, and the only detailed structural models currently available originate from computational approaches [19,32–34]. In addition, the length of the CLDN5 modulating peptides identified so far (30 amino acids for C5C2 and up to 170 amino acids for the modified cCPE) presents difficulties for both experimental and computational methods to study protein-peptide (P-Pept) interactions, not to speak about their potential translatability to the clinics. Indeed, while computational techniques can be crucial to understand the atomic details of the binding process, they are usually limited to peptides not exceeding 15 residues [35–40], with at least one remarkable and very recent exception [41]. As for the experimental side, shorter sequences are beneficial in terms of synthesis, solubility, conformational stability, and translatability [42].

In this work, we investigate a group of novel peptidomimetics extracted from either C5C2 or cCPE characterized by a short chain length (< 20 amino acids). Our study is based on a multidisciplinary approach, integrating computational methods and *in vitro* BBB models. Leveraging this workflow to screen a set of designed peptides, we identified a new 14-mer peptide (named *f1-C5C2*, sequence ESVLALNAEVQAAR, corresponding to the mCLDN5 fragment comprising the residues 68 to 81 and including the previously reported S74N substitution; [11]) that exhibits affinity for CLDN5 and increases the permeability in 2D murine *in vitro* models of BBB at relatively low concentrations and short incubation times. The multidisciplinary workflow described here is not specific to CLDN5, and hence can, in principle, be applied to other members of the CLDN family, providing new hints to design selective enhancers of drug delivery across different biological barriers and CLDN-based paracellular complexes.

## RESULTS AND DISCUSSION

### Design of short CLDN5-binding peptides

To predict whether the mCLDN5 peptidomimetics could compete with the interactions that stabilize the paracellular TJ scaffold, we visualized CLDN5 assemblies using structural models. Although there is no experimental structure available for CLDN-based TJ complexes, several models have been proposed [43]. Here, we used two CLDN5 multi-pore architectures, one based on the template previously published for the homolog mCLDN15 [43] (**Figure 1A-C**), and the other on the *back-to-back* interaction interface introduced for CLDN5 [32] (**Figure 1D-F**). Consistent with our previous work [34,44], these models were built using the human CLDN5 (hCLDN5) protomer. However, we emphasize that the sequence identity between the human (UNIPROT: O00501) and the murine (UNIPROT: O54942) CLDN5 sequences is ∼ 92%, as shown by the sequence alignment performed with Clustal [45,46] and reported in **Figure S1**. For this reason, the multi-pore configurations assembled using either hCLDN5 or mCLDN5 should not significantly differ.

**Figure 1.**
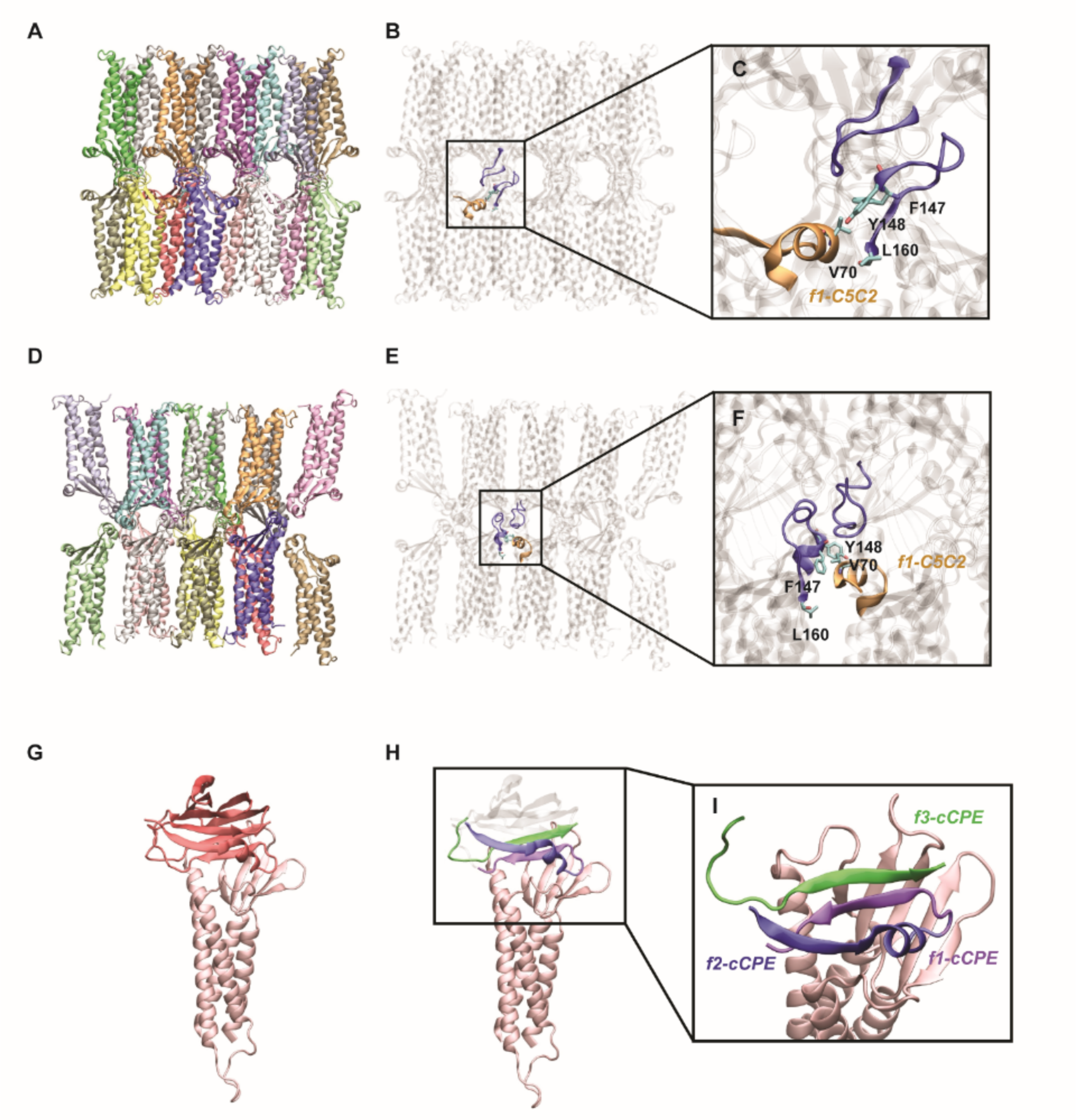
Selection of CLDN5- and cCPE-based peptides from structural models. **A.** Triple pore arrangement formed by multiple hCLDN5 monomers based on the mCLDN15 template [43]. Individual monomers are distinguished by color. **B.** Same structure as (A), with the f1-C5C2 fragment and the ECL2 segments shown in orange and blue, respectively. **C.** Close-up view of the V70, F147, Y148 and L160 residues forming the cis-linear interface. The ECL2 of the monomer belonging to the opposite hCLDN5 strand is also shown. **D.** Triple pore arrangement formed by multiple hCLDN5 monomers based on the CLDN-CLDN interface described in Ref. [32]. Individual monomers are distinguished by color. **E.** Same structure as in (D), with the f1-C5C2 fragment and the ECL2 segments shown in orange and blue, respectively. **F.** Close-up view of the V70, F147, Y148 and L160 residues forming the cis-linear interface. The ECL2 of the monomer belonging to the opposite hCLDN5 strand is also shown. **G.** hCLDN5-cCPE complex model generated with the modified version of the toxin [12] and aligned to the hCLDN4-cCPE crystal structure (PDB ID:5B2G). **H.** Same structure as in (G), in which the binding interfaces between hCLDN5 and cCPE are highlighted. **I.** Close-up view of the hCLDN5-cCPE contact domain, in which the cCPE fragments f1-cCPE, f2-cCPE and f3-cCPE are colored in purple, blue and green, respectively.

Short peptide sequences that could potentially bind CLDN5 were identified starting from sequences previously used to target this protein (C5C2 [11] and cCPE [12]). These oligomers, composed of more than 30 amino acids, were fragmented into shorter chains, with a maximal length of 16 residues. For the cCPE-derived ones, the domains responsible for the association with CLDN5 were considered [12]. In both human and murine architectures, the sequence of the first selected peptide, *f1-C5C2*, coincides with the region of the CLDN5 ECL1 domain responsible for the formation of both hydrophobic *cis-* and *trans*-interactions with adjacent and facing CLDN subunits, respectively (**Figure 1B,E**). Specifically, the *f1-C5C2* fragment represents the extracellular helix domain of CLDN5 protomers and includes the residue V70 (corresponding to M68 in mCLDN15) that stabilizes the so-called *cis-linear* interface [43] with the hydrophobic residues F147, Y148 and L160 (F146, F147 and L158 in mCLDN15) belonging to the adjacent monomer (**Figure 1C,F**). In addition, this segment is also involved in various *trans*-interactions that involve the apolar contact between L71 of the ECL1 of one monomer and residues P153 and V154 belonging to the ECL2 of the facing CLDN5 monomer. The second C5C2-based fragment, *f2-C5C2*, differs from *f1-C5C2* for the absence of the glutamate and the serine at the N-terminus, hence affecting its solubility in water.

cCPE-derived peptides were defined leveraging a model of the hCLDN5-cCPE complex. To assemble this system, a homology-based hCLDN5 protomer was aligned with the hCLDN4 crystal structure, which was solved bound to the bacterial toxin (PDB ID: 5B2G [29]). In addition, because CLDN5 is not a biological target for cCPE, the ligand was modified as suggested previously [12]. In the resulting hCLDN5-cCPE model (**Figure 1G**), three fragments (*f1-cCPE*, *f2-cCPE* and *f3-cCPE,* **Figure 1H,I**) with a length between 11 and 16 residues were selected from the binding interface. Thereafter, single-point mutations were introduced in the *f1-C5C2, f1-cCPE*, *f2-cCPE* and *f3-cCPE* sequences to improve their solubility in water, generating the peptide databank reported in **Table S1**.

### *In silico* and experimental assessment of peptide solubility

To predict the solubility of the peptide databank that encompasses both cCPE and C5C2-derived fragments, we performed all-atom MD simulations of multiple copies of each species in water and monitored the formation of aggregates. To properly classify each peptide, we designed a machine learning (ML) approach (**Figure S2)** based on three solubility descriptors calculated along the simulations: the number of peptides’ aggregates, the size of the largest aggregate, and the number of their contacts with water molecules (see **Section S1**). The method was trained on an external dataset of peptides of known solubility and allowed us to classify all our sequences into two neatly distinct families (see **Tables S1** and **S2** for test and train sets, respectively, and **Figure S3A**). We discuss here the results of the representative sub-group of peptides *f1-C5C2* and *f3-cCPE-mut1*, assigned to the soluble group, and *f2-C5C2*, included among the insoluble ones (**Figure S3B**). The presence of both soluble (*f1-C5C2*, *f3-cCPE-mut1*) and insoluble (*f2-C5C2*) molecules allowed us to assess the accuracy of the solubility prediction.

The starting and final configurations from MD simulations of each of the three peptides are shown in Figure 2 (panels **A,B; D,E;** and **G,H)**, while additional intermediate states along the trajectories are reported in **Figure S4**. During the last 10 ns of the simulation, we observed that the soluble *f1-C5C2* and *f3-cCPE-mut1* peptides form only a few aggregates with an average size of up to ∼ 7 and ∼ 10 subunits, respectively (∼ 30% and ∼ 37% of the number of initial copies for *f1-C5C2* and *f3-cCPEmut1,* respectively). In addition, these oligomers were significantly dispersed in the solvent, as demonstrated by the presence of ∼11 different aggregates at the end of the simulation for both systems (**Table S1**). As for the extent of solvation, *f1-C5C2* and *f3-cCPE-mut1* maintained more than 75% contact with water molecules after 100 ns of MD simulation. On the contrary, *f2-C5C2* formed a cluster of 12 subunits, about 45% of the total copies, and only four large aggregates were found at the end of the simulation, suggesting a marked propensity of the oligomers to interact with each other, rather than being individually solvated. Indeed, at the end of the simulation, the number of contacts with water molecules for *f2-C5C2* was ∼ 65% of the initial value. This suggests that the surface exposed to the solvent for the system assembled with *f2-C5C2* is significantly lower than those of *f1-C5C2* and *f3-cCPE-mut1.* Remarkably, liquid chromatography-coupled mass spectrometry (LC-MS) fully confirmed the computational predictions on solubility, as illustrated in Figure 2 (panels **C,F,I**).

**Figure 2.**
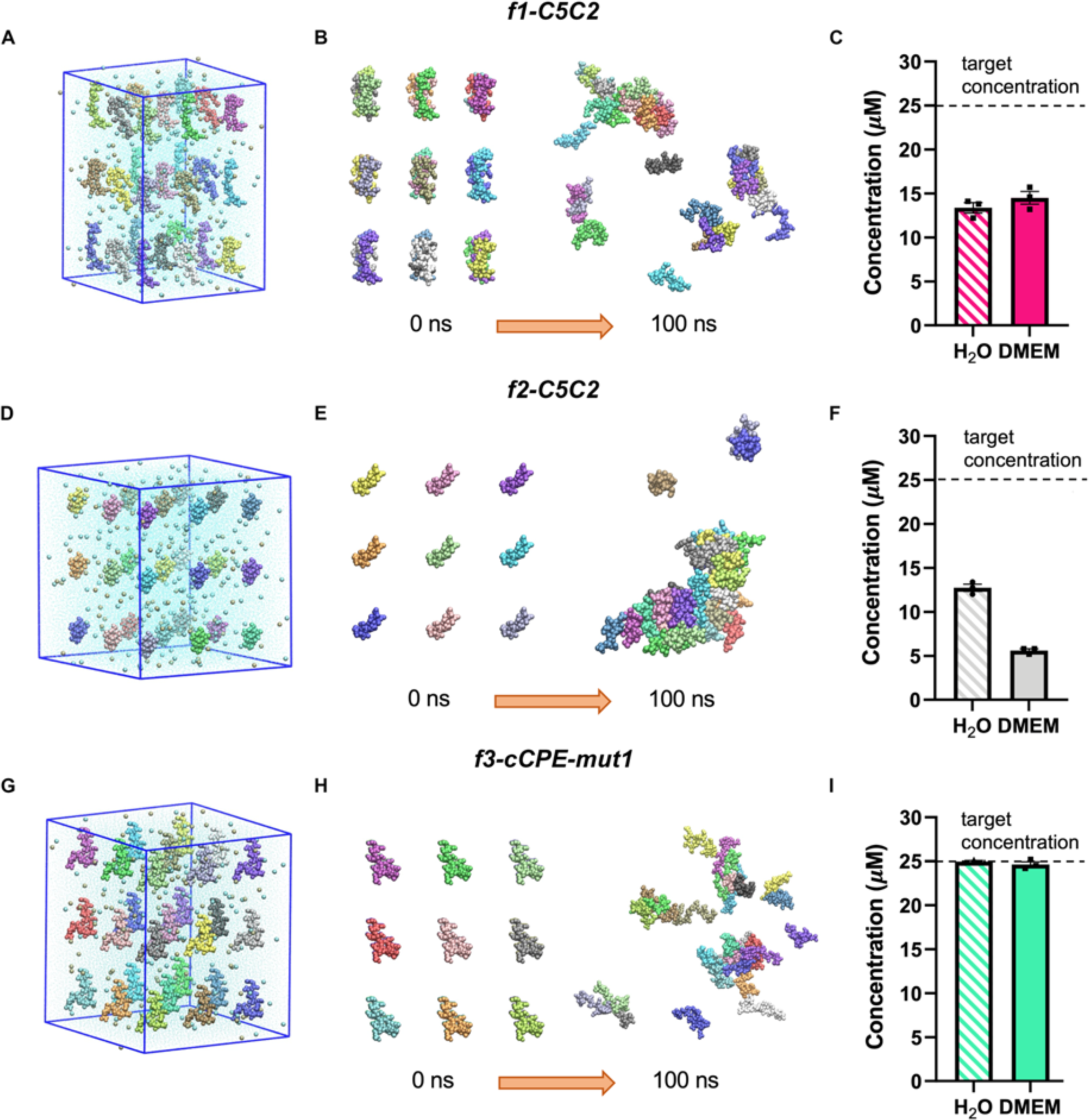
Computational and experimental assessment of the peptide water solubility. **A,D,G.** Rectangular simulation boxes composed of 27 identical peptides arranged in a three-dimensional grid, solvated with water and the mixed KCl/CaCl2 ionic bath for f1-C5C2 (**A**), f2-C5C2 (**D**), and f3-cCPE-mut1 (**G**). The volumes of each simulation box is related to the size of the specific peptide. **B,E,H.** Peptide configuration at the beginning and at the end of the 100-ns-long MD simulation for f1-C5C2 (**B**), f2-C5C2 (**E**), and f3-cCPE-mut1 (**H**). **C,F,I.** Experimentally determined peptide solubility measured by LC-MS for f1-C5C2 (**C**), f2-C5C2 (**F**), and f3-cCPE-mut1 (**I**). For LC-MS experiments, peptides were dissolved in H2O:AcN (1:1), and different concentrations were used for the calibration curve. The three samples were then diluted in H2O and culture medium (DMEM) to a final concentration of 25 μM (working concentration in vitro). The graphs report the absolute concentration found by LC-MS analysis.

### Standard MD simulations assess the stability of mCLDN5-peptide complexes

After the solubility assessment, we proceeded with studying the binding of *f1-C5C2* and *f3-cCPE-mut1* to mCLDN5. We first modeled the peptide-protein complexes by performing molecular docking calculations where the flexibility of the peptides was taken into account using an iterative procedure to scan all torsion angles [47–49]. Then, the resulting docked conformations were subjected to 200 ns of standard, all-atom MD simulations, restricting the volume accessible to the peptide within the cylindrical domain shown in **Figure S5**. The trajectories can be visualized in the **Video S1**.

For mCLDN5-*f1-C5C2*, we observed that the complex maintains a bound state, although with a significant rearrangement of the peptide, which moves towards the mCLDN5 ECL1 domain (Figure 3A). On the contrary, the mCLDN5-*f3-cCPE-mut* complex dissociates over the same time scale (Figure 3B), with the peptide diffusing towards the solvent. Hence, these preliminary simulations already suggest a different affinity of the two peptides for mCLDN5.

**Figure 3.**
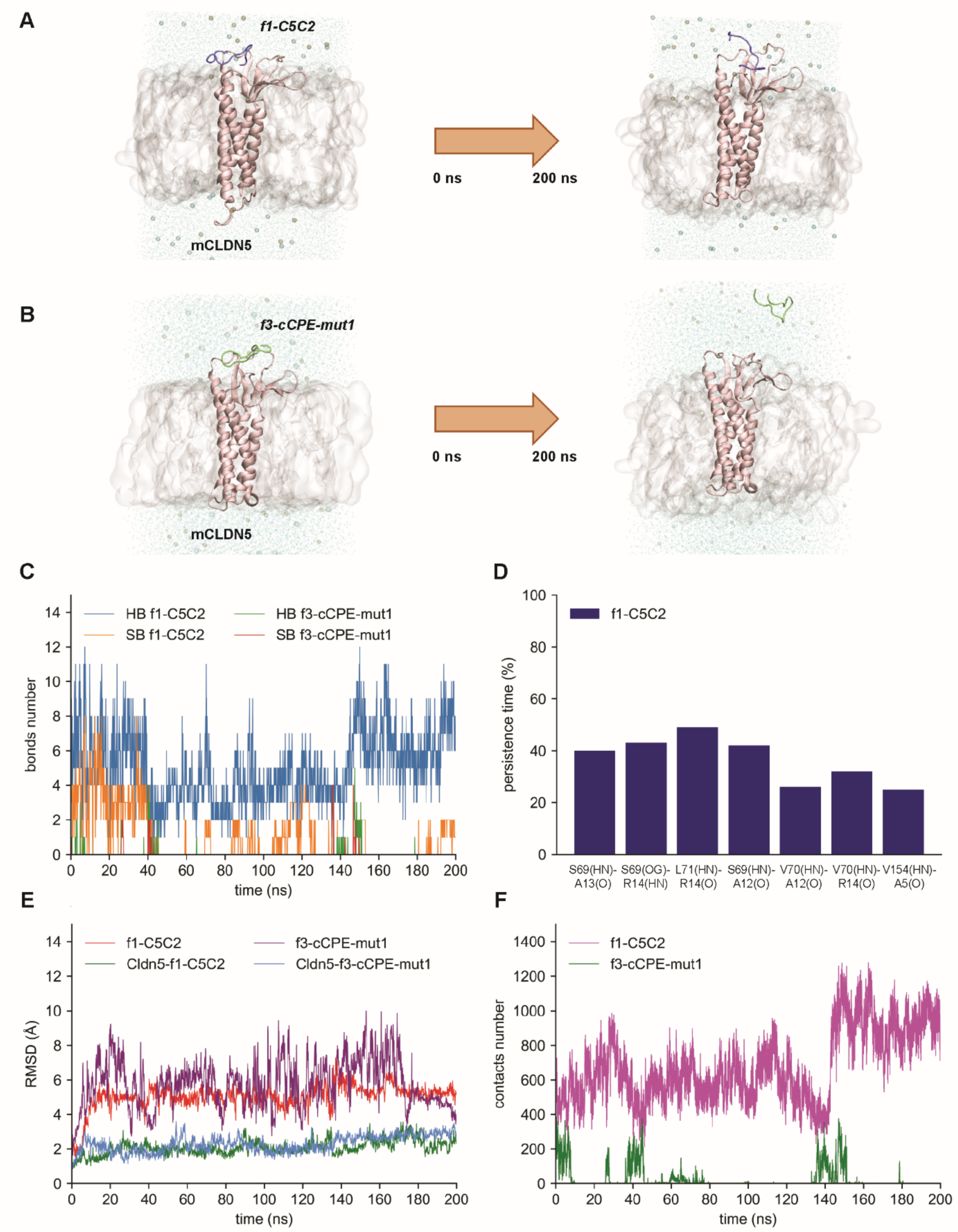
Standard MD simulations of mCLDN5-peptide complexes. **A,B.** Initial and final snapshots of the simulated trajectories of mCLDN5-f1-C5C2 (**A**) and mCLDN5-f3-cCPE-mut1 (**B**) complexes surrounded by the membrane bilayer (grey solid surface), water molecules (cyan points) and KCl ionic bath (blue and yellow transparent VdW spheres for K^+^ and Cl^-^, respectively). mCLDN5 is shown with a pink cartoon representation. The f1-C5C2 and f3-cCPE-mut1 peptides are indicated as blue and green unfolded coils, respectively. **C.** Number of hydrogen bonds (HBs) and salt bridges (SBs) formed between the peptide and the mCLDN5 ECL domain for f1-C5C2 and f3-cCPE-mut1. **D.** HBs with a persistence time > 20% of the total simulated time for f1-C5C2 are shown as blue bars. No HBs with a persistence > 20 % of the total simulated time were found for f3-cCPE-mut1. **E.** Backbone RMSD of the peptide and the mCLDN5 ECL domain. **F.** Contact number between the mCLDN5 ECL and the peptide during the MD simulation.

For both systems we calculated the time evolution of the number of hydrogen bonds (HBs) and salt bridges (SBs) between protein and peptides, the HB persistence time, the root-mean square deviation (RMSD) of the peptide, the RMSD of the mCLDN5 ECL domain, and the contact number between the protein and the peptide. The changes in the number of HBs/SBs in the mCLDN5-*f1-C5C2* and mCLDN5-*f3-cCPE-mut1* complexes (Figure 3C**,D**) quantify the different interactions of the two ligands with mCLDN5: while *f1-C5C2* formed various HB with a persistence time > 20 % (Figure 3D) involving residues S69, V70, L71 and V154 of mCLDN5 (**Figure S6**), no persistent interactions were found for *f3-cCPE-mut1*. The RMSD values of the mCLDN5 ECL domain remained stable during the MD simulations of both systems (green and cyan plots, Figure 3E), while the RMSDs of the peptide backbone were subjected to remarkable modifications. At about ∼ 20 ns, *f1-C5C2* switched to a novel stable configuration with an RMSD of ∼ 5 Å (red plot, Figure 3E), while the *f3-cCPE-mut1* conformation was greatly changing, with the RMSD reaching up to ∼ 9 Å (purple plot, Figure 3E), as a result of the increasing exposure to the solvent. Finally, the number of contacts between the protein and *f1-C5C2* rose up to ∼ 1100, while the same quantity was negligible in the case of the second system (Figure 3F).

### Free Energy calculations reveal the binding mechanism between *f1-C5C2* and mCLDN5

We used the Temperature Accelerated Molecular Dynamics (TAMD) approach [50,51] to study the unbinding of *f1-C5C2* and *f3-cCPE-mut1* from mCLDN5, using the distance between the centers of mass (COM) of the mCLDN5 ECL domain and the COM of each peptide as a collective variable (CV). The FE profiles of each protein-peptide complex are shown in Figure 4A. The curve for *f1-C5C2* is characterized by a minimum of ∼ -6 kcal/mol at the COM-COM distance of ∼ 10.3 Å. Beyond this distance, FE increases until it reaches a plateau around 17.0 Å, stabilizing at a value of 0 kcal/mol, corresponding to the peptide in the bulk solvent. Using the FE profile of the complex and the formalism described in the Material and Methods, we estimate an equilibrium dissociation constant (Kd) between mCLDN5 and *f1-C5C2* of 57.9 µM. On the contrary, the *f3-cCPE-mut1* FE profile shows a high energy barrier with a maximum value of ∼ 9 kcal/mol at the smallest COM-COM distance sampled (8 Å). As the distance increases, FE gradually decreases up to a distance of ∼ 17 Å, where it reaches a value of 0 kcal/mol that is maintained with minimal fluctuations up to the COM-COM distance of 30.0 Å. Taken together, these results confirm the binding of *f1-C5C2* to mCLDN5 and the repulsive interaction of the cCPE-derived peptide with the same protein. As a validation test, we repeated the FE calculations using the extended-system adaptive biasing force (eABF) method [52] and obtained similar results (see **Section S2** and **Figure S7)**. A comparative analysis of the convergence rates of the two approaches is reported in **Figure S8**.

**Figure 4.**
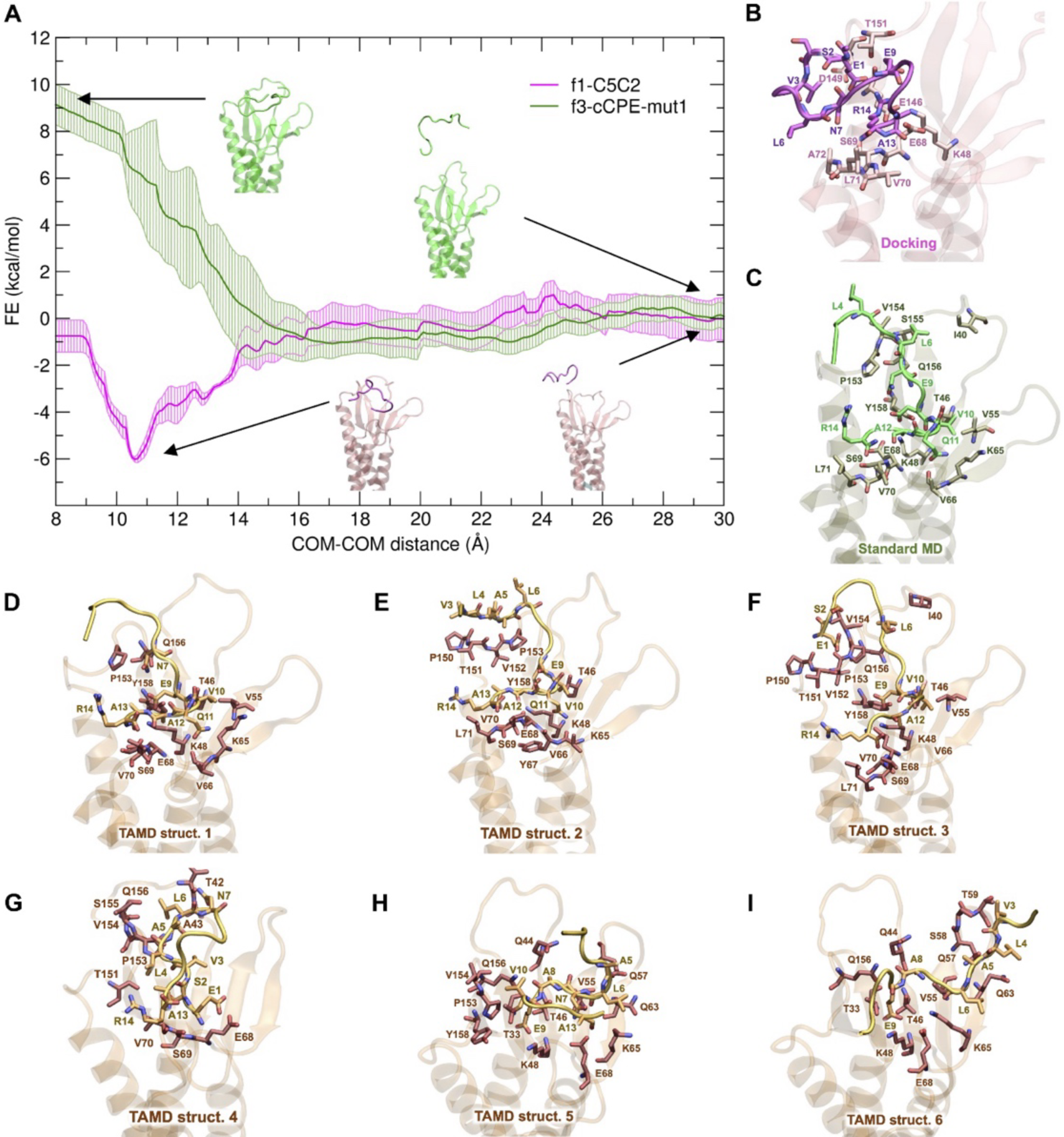
mCLDN5-peptide unbinding free energy calculations using TAMD. **A.** Free energy (FE) profiles for f1-C5C2 (magenta) and f3-cCPE-mut1 (green) are computed with respect to the mCLDN5-peptide COM-COM distance with the TAMD algorithm. FE values and errors are indicated as the mean and standard deviation from three 150 ns-long replicates. Representative structures of the CLDN5-peptide complex are shown and colored in magenta and green for f1-C5C2 and f3-cCPE-mut1, respectively. **B-I.** Representative mCLDN5-f1-C5C2 binding configurations provided by docking, standard MD and clustering of TAMD trajectories. Amino acids involved in the interactions are shown as sticks and listed in **Table S3.**

The enhanced simulations allowed us to improve the sampling of the mCLDN5-*f1-C5C2* poses with respect to molecular docking and standard MD simulations (whose resulting conformations are shown in Figure 4B and **4C**, respectively). From an RMSD-based clustering of the TAMD runs, we obtained six representative structures (Figure 4D-I) that we analyzed for residue-residue interactions between mCLDN5 and the peptide (**Table S3**). Interestingly, most of the contacts involve the protein domains, including residues E68, S69, V70, and L71 of ECL1, and T151, P153, V154, and Y158 of ECL2, analogously to what was observed using the hCLDN5 triple-pore models (see Figure 1). The same analysis was performed on the eABF trajectories, obtaining very similar interaction networks, as shown in **Figure S7D-I**.

### Experimental assessment of the mCLDN5-peptide binding affinity

The binding affinity of selected peptides for mCLDN5 was estimated using microscale thermophoresis (MST) [53,54], which allows calculating the Kd of a binding reaction by measuring variations in protein fluorescence intensity caused by a ligand (peptide) binding through a laser-induced microscopic temperature gradient. To obtain fluorescently-labeled mCLDN5 while maintaining the structural integrity of the transmembrane complex, we transfected HEK-293T cells, that physiologically do not express CLDN5 [55], with a tGFP– tagged mCLDN5 construct (Figure 5A). We then employed cell lysates for the following MST measurements. On average, GFP (and, therefore, mCLDN5) represented 1/250 of the total protein content.

**Figure 5.**
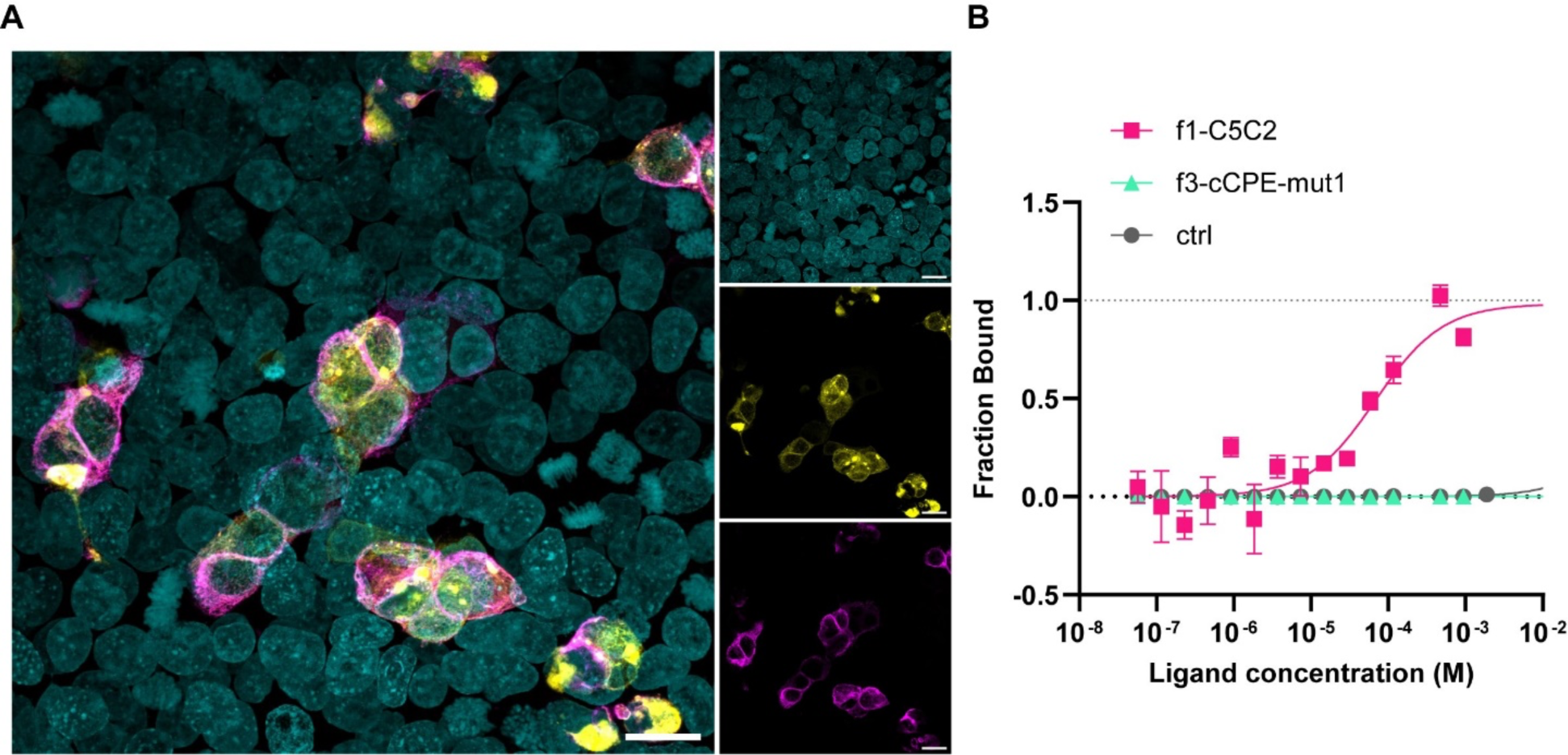
Assessment of the mCLDN5-peptide binding affinity. **A.** Representative confocal imaging of HEK-293T cells transfected with a tGFP-tagged mCLDN5 construct and subjected to immunocytochemistry with mCLDN5 antibodies. Nuclei are in cyan, GFP in yellow, mCLDN5 in pink. Scale bars, 20 µm. **B.** MST binding curves and extracted Kd for f1-C5C2 (red symbols) and f3-cCPE-mut1 (green symbols) peptides, together with a negative control sample (black symbols; recombinant tGFP spotted in the cell lysate in the absence of transfected mCLDN5). The extracted Kd value for f1-C5C2 was 68.26 µM.

To determine the Kd of peptide binding, HEK-293T lysates containing GFP-mCLDN5 (1 μM) were incubated in MST capillaries with increasing concentrations of soluble *f1-C5C2* and *f3-cCPE-mut1* peptides. From the analysis shown in Figure 5B, the resulting Kd of *f1-C5C2* was ∼ 68 μM, in excellent agreement with the value estimated from TAMD simulations (∼ 58 μM), while experiments with *f3-cCPE-mut1* and control did not display significant binding affinity. These results fully confirmed the different peptides’ selectivity for mCLDN5 proteins obtained from the FE calculations.

### Peptide-mediated opening of paracellular spaces in brain endothelial cells

We then moved to investigate the functional effects of the selected peptides on an *in vitro* murine 2D transwell model of BBB. The model was validated by monitoring the trans-endothelial electrical resistance (TEER) of the cell layer and the expression of mCLDN5, as shown in our previous work [56] and in **Figure S9**. All experiments were performed when the TEER values reached a plateau, indicating the correct formation of the TJs and the integrity of the endothelial barrier. The selected peptides, at the working concentration of 25 μM, did not affect the viability of the endothelial cell layer, as assessed using a live/dead assay (**Figure S10**). We then monitored the changes in TEER values upon incubation with 25 μM of either soluble peptides *f1-C5C2* and *f3-cCPE-mut1* (Figure 6A-C). We observed an immediate and progressive TEER decrease for cells incubated with *f1-C5C2,* becoming highly significant after 8 h of treatment. The decrease in TEER values (around 40% of the baseline) can be considered a proxy for the partial opening of paracellular spaces. On the contrary, TEER values remained constant when *f3-cCPE-mut1* was used, in line with the results obtained for FE calculation and binding affinity. The working concentration was established from the preliminary solubility check with MS (see Figure 2) and represents, to the best of our knowledge, the lower peptide concentration used so far for this purpose [11,12,14].

**Figure 6.**
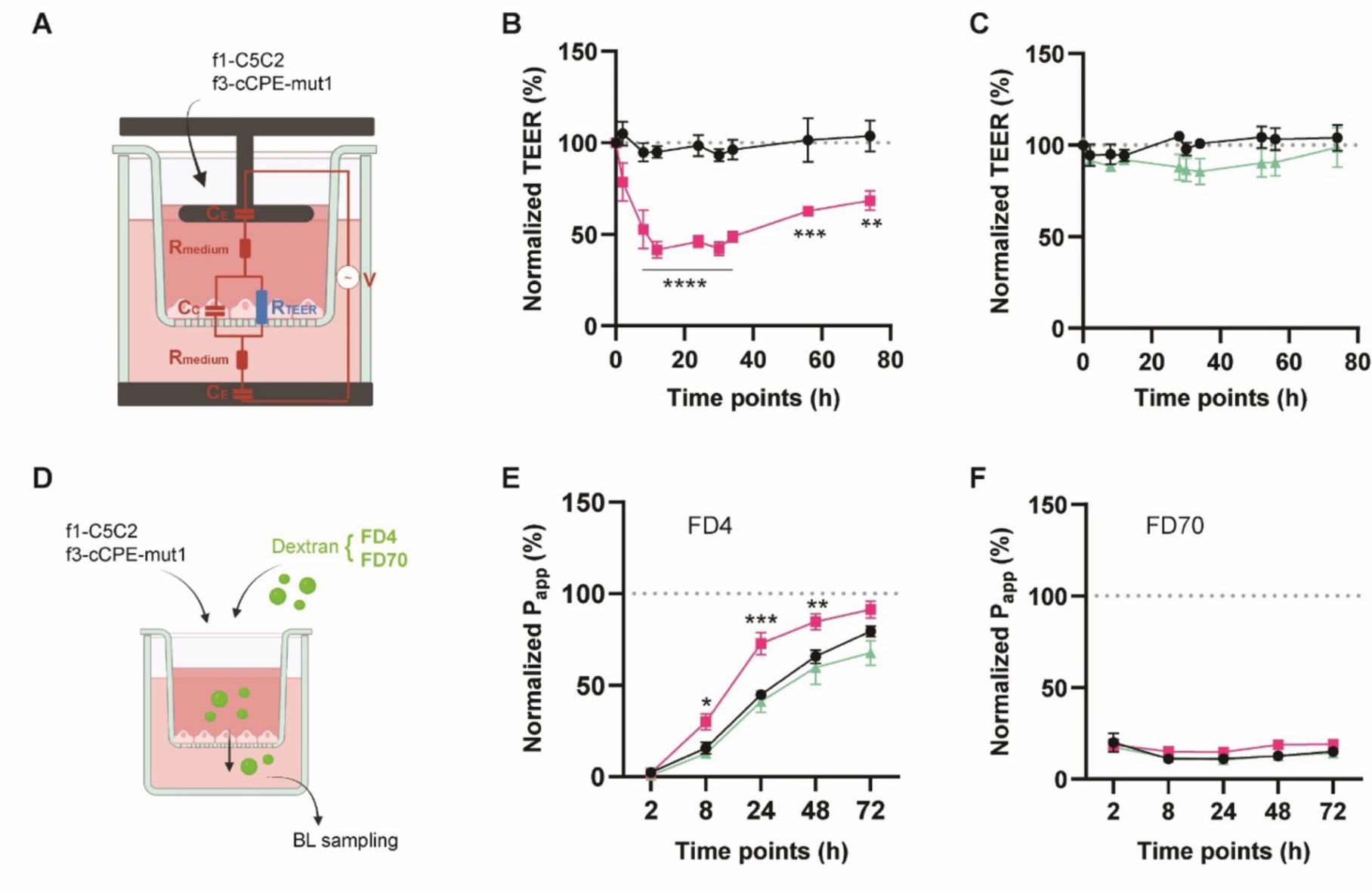
Effect of the exposure to selected peptides on the in vitro BBB properties. **A.C.** Schematic representation of TEER experiments (**A**). TEER values (expressed as percentages of baseline values) for bEnd.3 cell layers exposed to f1-C5C2 (**B**) and f3-cCPE-mut1 (**C**). **D.** Schematic representation of 4/70 kDa fluorescein isothiocyanate-dextran transport studies. **E.** Normalized apparent permeability of FD4 (expressed in percent of the permeability in the absence of cells) for bEnd.3 cell layers exposed to either f1-C5C2 (pink square) or f3-cCPE-mut1 (green triangle). **F.** Normalized apparent permeability of FD70 (expressed as in percent of the permeability in the absence of cells) for bEnd.3 cell layers exposed to either f1-C5C2 (pink square) or f3-cCPE-mut1 (green triangle). Working concentrations for both peptides were 25 μM. Data are means ± sem (n = 3), *p < 0.1, **p < 0.01, ***p < 0.001, ****p < 0.0001 one-way and two-ways ANOVA/Tukey’s tests versus control conditions (vehicle).

As a complementary assay, we also assessed the capability of fluorescein-labeled dextran of different sizes (4 KDa for FD4 and 70 KDa for FD70) to translocate across the barrier (Figure 6D-F). Notably, FD4 is a well-known hydrophylic marker for paracellular transport [57,58]. We incubated each molecule in the presence of of either *f1-C5C2* or *f3-cCPE-mut1* (25 μM) and monitored their apparent permeability (Papp, see more details in the Materials and Methods) at various time points from 2 to 72 h of incubation (see scheme in Figure 6D).

Data were expressed in percent of the maximum permeability that was determined in the absence of cells (empty transwells), where FD4 and FD70 freely and equally distribute into the apical to the basolateral compartments across the semipermeable membrane down their concentration gradient. The progressive increase in the FD4 translocation through the cell barrier over time under control conditions (vehicle) indicates the presence of transcellular transport of the low molecular mass fluorescent tracer, while FD70 was not significantly transported (Figure 6E**,F**). Interestingly, *f1-C5C2* showed a significant increase in the kinetics of the Papp of FD4 starting from 8 h of incubation (Figure 6E**,F**), in good agreement with the observed decrease in TEER in the same time window. On the contrary, the Papp of the larger FD70 was not affected by exposure to the *f1-C5C2* peptide (Figure 6F). This indicates that the opening of the paracellular spaces by the *f1-C5C2* peptide exhibit size selectivity that represents a crucial aspect to avoid an indiscriminate translocation of any species (including pathogens) to the brain during the paracellular opening window. For both FD4 and FD70, the transport behavior did not differ from the control conditions when *f3-cCPE-mut1* was used (Figure 6E**,F**).

### Subcellular distribution of mCLDN5 upon peptide-mediated opening of paracellular spaces

mCLDN5 expression is linked to that of other TJ proteins, and all have a strict relation with the submembranous actin cytoskeleton [59–61]. Thus, we quantified the total expression of mCLDN5, Occludin *(OCLN),* and Zonula Occludens 1 (ZO1) through Immunoblotting upon exposure to 10 and 25 µM of *f1-C5C2* for 24 h (Figure 7A-D). Both transmembrane proteins (mCLDN5 and OCLN) and the cytoplasmic scaffolding protein ZO-1 were significantly downregulated when 25 µM *f1-C5C2* was used, while actin remained unaltered.

**Figure 7.**
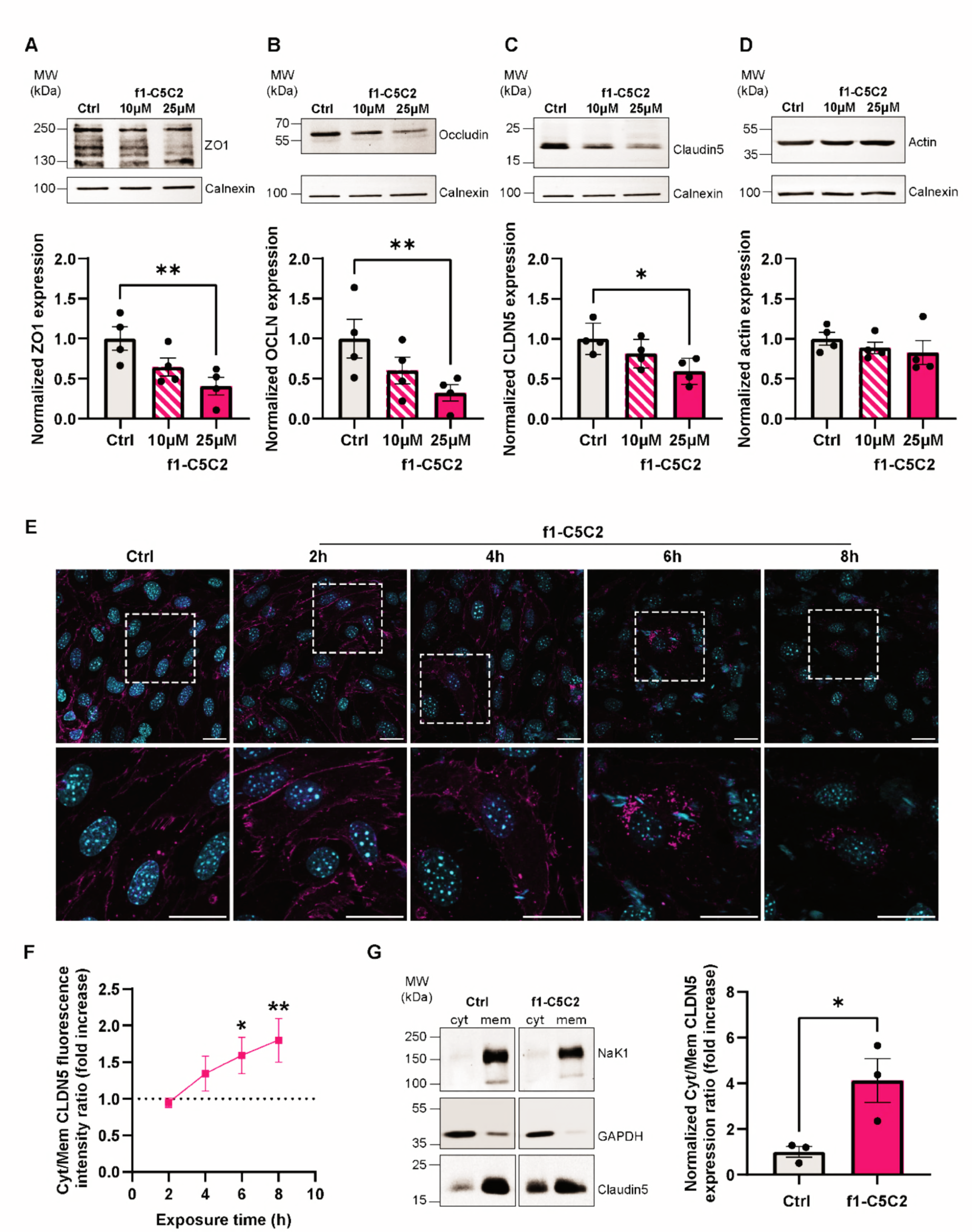
Expression and translocation of TJ proteins upon exposure of bEnd.3 cells to the f1-C5C2 peptide. **A-D.** Upper panels: Representative Western blots stained with ZO1 (220 kDa), OCLN (55kDa), CLDN5 (23 kDa), Actin (42 kDa) and calnexin (100 kDa) for bEnd.3 cells treated with vehicle (Ctrl) or exposed to 10 and 25 μM f1-C5C2. Lower panels: Quantification of protein expression normalized over the respective controls. Data are means ± sem (n = 4), *p < 0.05, **p < 0.01, one-way ANOVA/Friedman’s test. **E.** Representative confocal images for mCLDN5 immunostaining at various time points (2, 4, 6 and 8 h) for bEnd.3 cells treated with vehicle (Ctrl) or exposed to 10 and 25 μM f1-C5C2. The boxed areas are shown under each image at higher magnification. Scale bars, 25 μm. **F.** Results of the quantitative image analysis mCLDN5 subcellular distribution. Values of fluorescence intensity ratio of cytosolic and plasma membrane CLDN5 at various time points (2, 4, 6 and 8 h) were normalized over the respective controls. Data are means ± sem (n = 3), *p < 0.05, ** p < 0.01, ordinary one-way ANOVA test. **G.** Surface biotinylation of bEnd.3 cells treated with vehicle (Ctrl) or exposed to 25 μM f1-C5C2. Left panel: Representative blots stained with antibodies to mCLDN5 (18 kDa), the intracellular housekeeping protein GAPDH (36 kDa), and the plasma membrane-associated Na^+^/K^+^-ATPase 1 (NaK1; 130 KDa). Right panel: Quantification of mCLDN5 intracellular (cytosolic)/extracellular (membrane) expression normalized over the respective controls. Data are means ± sem (n = 3), *p < 0.05, unpaired Student’s t-test.

We then carried out immunofluorescence experiments to follow the subcellular trafficking of immunostained mCLDN5 over time following exposure to the *f1-C5C2* peptide (Figure 7E). Quantitative confocal imaging analysis reported in Figure 7F (see **Figure S11** for more details) showed that the cytosolic/membrane ratio of mCLDN5 in bEnd.3 cells increased significantly after 6-8 h of incubation with *f1-C5C2* peptide, indicating a marked internalization of the protein, concomitant with the TEER lowering shown in Figure 6A. This result was fully confirmed by surface biotinylation experiments in live bEnd.3 cells treated with either vehicle or the *f1-C5C2* peptide for 24 h, which showed that the biotinylated mCLDN5 fraction exposed on the plasma membrane was significantly decreased in favor of its cytosolic fraction that was correspondingly increased, in the absence of changes in the cytosolic and plasma membrane markers GAPDH and Na^+^/K^+^-ATPase-1, respectively (Figure 7G).

## CONCLUSIONS

The treatment of many neurological pathologies and the development of novel drugs against CNS-related disorders are still hampered by the limited delivery across the BBB. In this context, many strategies are under investigation to improve the passage of small molecules across the BBB, by enhancing either the transcellular or the paracellular pathways. The investigation of possible TJ-modulating strategies to enable small molecular weight drugs or nutrients to cross the BBB through the paracellular route is gaining increasing interest for its translation application in a variety of genetic or pathological brain conditions. Within BBB TJs, CLDN5 proteins are essential in sealing the paracellular space between two adjacent cells, and they could act as receptors to bind TJ-modulators in order to enhance BBB permeability in a safe and reversible manner. TJ-modulators hold clear advantages compared, for instance, to absorption enhancers (e.g., sodium caprate, chitosan, or alkylglycerols) [62,63] as they can specifically and directly interact with structural components of the TJs. Direct TJ modulators are mainly classified into two types: RNA interference agents and binding molecules (binders), which include derivatives of the non-toxic C-terminal domain of *Clostridium perfringens* enterotoxin (cCPE) [12,26,27], peptidomimetics based on the murine CLDN5 (mCLDN5) ECL1 domain [11,27] and, in some cases, antibodies [64,65]. Peptide-based TJ modulators offer the opportunity of a widespread (opposite to localized approaches such as ultrasound Focused ultrasound (FUS)-mediated BBB opening [66]) and transient action (given their short half-life in biological environments [67]), and also allow for flexible engineering.

In this work, we exploited this opportunity setting-up a synergistic *in silico-in vitro* platform, allowing for screening CLDN5-binding peptides presenting suitable solubility in biological media and efficient protein binding. We identified a short peptide of 14 amino acids, here called *f1*-C5C2 and demonstrated that is is a valuable CLDN5-derived candidate for transient BBB permeabilization. The reduced size of the peptide constitutes a big advantage in terms of solubility and effective concentration to be used. Our data have shown that the peptide associates with the CLDN5extracellular domain [11,14,68], recapitulating, for instance, the binding of cCPE derivatives to mCLDN5 [12] and to other crystallized claudins [29–31,69]. Given the very short half-life of CLDN5 oligomers, the role of the peptides would be to prevent the recruitment of new CLDN5 monomers to the paracellular complex, thereby increasing paracellular delivery across the BBB. Previous reports are consistent with our observation that, following the tetramer complex opening, the mCLDN5-peptide complexes can be endocytosed by brain endothelial cells [70–72]. This mechanism would alter the distribution of mCLDN5 proteins at the membrane (cell-cell contact), thereby further impairing the integrity of TJ strands. This is a very important aspect that deserves further investigation in future works. Indeed, while mislocalization of occludin might initiate caspase activation [73], it has also been shown that downregulation of total mCLDN5 expression by binding peptides might have beneficial effects in other pathological contexts, as previously reported for alcohol-fed rats [74]. CLDN5 knockout strategies in mice, associated with increased paracellular transport of small molecules (800-1,000 Da), showed no TJ breakdown or edema formation. Rather, after traumatic brain injury, CLDN5 knockout was associated with a reduced focal cerebral edema and improved cognitive functions [75–77]. Understanding the precise implication of TJ dysregulation will be of pivotal importance to finely control this BBB opening tools.

In conclusion, we set up a robust *in silico*/experimental workflow for the design and validation of competitive peptides to open CLDN5 multimeric assemblies. Our platform, which includes complementary validation of peptide solubility, binding affinity, and paracellular space opening, can be widely applied to screen potential short peptides with a potential affinity for any TJ protein. The use of efficient short peptides would also be very beneficial in simplifying the synthetic process, increasing the solubility, and lowering the doses. We have shown the capability of a short C5C2-derived peptide to target and bind mCLDN5 complexes with high specificity, inducing lowering of TEER values and increased paracellular permeability to small molecules, in a size-selective manner at very low concentrations and short incubation times. To convert these findings into feasible clinical applications, additional efforts are needed to fully understand and control the mechanism of action and the reversibility of the process. Despite a long way to go, we believe that our work sets a solid basis for the development of this promising field.

## MATERIALS AND METHODS

### Design of the peptide dataset

Starting from structural models of the mCLDN5-cCPE and multi-pore hCLDN5 complexes, we generated shorter fragments of either the bacterial toxin cCPE or the peptidomimetic C5C2 [11] by removing the segments not directly interacting with CLDN5. We obtained three peptides from cCPE (*f1-cCPE, f2-cCPE, f3-cCPE*) and two from C5C2 (*f1-C5C2, f2-C5C2*). Then, we introduced single-point mutations in the resulting sequences to enhance their hydrophilicity, generating the peptides databank shown in **Table S1**. A detailed description of the procedure adopted to design each oligomer is provided in **Section S3**, whereas the generation of hCLDN5 triple-pore models, based on the strategy used in our previous works [34,44,78–80], is illustrated in **Section S4**.

### Peptide structure prediction, equilibration, and computational assessment of water solubility

After *de novo* modeling and preliminary relaxation, each peptide was solvated and equilibrated for 100 ns by standard MD simulations in the NPT ensemble at T=303.15 K and P=1 bar maintained by a Langevin thermostat and Nosé-Hoover Langevin barostat [81,82]. Simulations were performed with the CHARMM36m force field [83] and the NAMD 3.0 program [84]. More details on the procedure are reported in **Section S5**. To predict the propensity the peptides listed in **Table S1** to either form aggregates or remain adequately dispersed when immersed in water, we devised a computational strategy based on the introduction of descriptors for molecular aggregates formed in the simulations, as described in prior work [85]. The full protocol is sketched in **Figure S2** and discussed in **Section S6**.

### MD simulations of the mCLDN5-peptide complexes

Because of the absence of an experimental template, the mCLDN5 monomeric structure was generated by homology modeling using either the mCLDN15 (PDB ID: 4P79 [28]) or the hCLDN4 (PDB ID: 5B2G) crystal structures as templates. After the refinement of the model via all-atom MD simulations (see **Section S7**), the equilibrated configuration of the protein was used as a receptor to produce an initial pose for the mCLDN5-peptide complexes with *f1-C5C2* and *f3-cCPE-mut1* using molecular docking clalculations. Since the number of dihedral angles in our ligands is larger than what allowed in the docking program, we employed an iterative procedure based on the optimization of poses for adjacent peptide segments. Details are reported in **Section S8**.

Selected models for the mCLDN5-*f1-C5C2* and the mCLDN5-*f3-cCPE-mut1* systems were then embedded in a 1-palmitoyl-2-oleoyl-sn-glycero-3-phosphocholine (POPC) lipid bilayer, solvated with water and a 0.15 M KCl ionic bath. After a preliminary equilibration of 30 ns, 200 ns of MD simulations were performed with NAMD 3.0 [84] and the CHARMM36m force field [86] in the NPT ensemble at T=310 K and P=1 bar maintained by a Langevin thermostat and the Nosé-Hoover Langevin piston pressure control [81,82]. During production, the protein was restrained from lateral movement, and the peptide rearrangement was confined in a cylindrical region to prevent lateral diffusion, as shown in **Figure S5**. The two MD trajectories were analyzed with VMD 1.9.3 [87] for structural descriptors and can be visualized in the **Video S1**. Additional details of the standard MD simulations and the analysis are provided in **Section S9**.

### Claudin-peptides unbinding MD simulations

Enhanced sampling approaches can significantly increase the efficiency of MD simulations, in particular for highly flexible systems such as protein-peptide complexes [48,49,88–91]. Several techniques can be used to speed up the conformational sampling of the peptide and accelerate its escape from the energy minimum in the binding site, rarely seen in standard MD simulations [90]. Some of the options include applying a higher temperature to selected degrees of freedom [50,92], using constant [93] or adaptive [92] biasing potentials, or canceling the mean force acting on the collective variables (CVs) [94,95]. Here, free energy (FE) profiles for peptide unbinding from mCLDN5 were calculated using Temperature Accelerated MD (TAMD) simulations [50,51], taking as CV the distance between the centers of mass (COMs) of the peptides and the mCLDN5’s ECL, mapped in a range from 8 to 30 Å. The lateral diffusion of the peptide was limited by a cylindrical restraint, as done already in standard MD simulations (**Figure S2**). The use of spatial restraints was recommended in previous works [88,96–100] to ensure a faster convergence of the calculations, and their effect must be taken into account when computing the dissociation constant (see below). While TAMD is an efficient method to enhance sampling [92,101], other techniques have been developed to obtain the FE from a TAMD trajectory by integrating the mean forces acting on each CV (i.e. minus the derivative of the FE), such as the single-sweep (SSw) [102,103] and the on-the-fly parametrization (OTFP) methods [104,105]. Here, TAMD was performed using the Colvars module [106], via the extended Lagrangian dynamics feature for the CV [107], and the FE was computed from the mean forces collected along the runs [52]. More details on the approach and its implementation are reported in **Section S10**. To validate the FE profiles obtained from TAMD, we calculated them also using the extended-system adaptive biasing force (eABF) approach [52,94]. Finally, the mCLDN5-*f1-C5C2* FE curve from TAMD was used to predict the dissociation constant, Kd. Further information on eABF and the calculation of Kd are provided in **Section S10**.

### Clustering structures

Representative structures of the mCLDN5-peptide complex obtained from TAMD simulations were selected by filtering the configurations showing a distance between the peptide and the mCLDN5 extracellular COMs corresponding to the FE minimum ± 0.5 Å. The resulting structures were grouped in clusters based on the RMSD of the peptides, calculated with respect to the final configuration resulting from the standard MD simulation. In each cluster, a threshold of 2.5 Å was chosen, thus generating six ensembles ranging from 0 to 15 Å. The center of each cluster was selected as a representative structure of the sampling performed during the TAMD simulations. The same clustering procedure was performed for the eABF structures, although they were not used for further investigations.

### Peptide synthesis and mass spectrometry analysis of peptide solubility

All peptides were custom synthesized by Proteogenix. A 7.5 mM stock solution in H2O/ACN 1:1 was prepared and kept at -20 °C. For experiments with cells, dilutions from the stock were prepared in fresh cell culture medium and sonicated for 5 min in an ultrasonic bath. H2O/ACN 1:1 at the same dilution was used as a control. *f1-C5C2*, *f2-C5C2*, *f3-cCPE-mut1* peptides were dissolved in milliQ water for the preparation of 1 mM stock solution for each peptide. For an external calibration, 8 standard solutions for each peptide were prepared in H2O/ACN 1:1 (10 pM, 100 pM, 1 nM, 10 nM, 100 nM, 1 µM, 10 µM and 30 µM). Each peptide was then diluted in milliQ water, Dulbecco’s Modified Eagle Medium (DMEM) and cDMEM to reach a concentration of 25 μM and left for 1 h at 37 °C prior Mass Spectrometry analysis. All data were acquired on Waters® ACQUITY® SQ Detector, coupled with Waters Acquity UPLC system. Peptides were loaded on Acquity UPLC BEH C18 column (2.1 mm x 50 mm, 1,7 μm particle size). Ionization was conducted by positive ESI mode, the capillary voltage was set to 2.5 kV and the cone voltage to 35 V. Selected Ion Recording (SIR) of two masses ([M+3H]^3+^ = 606.1; [M+2H]^2+^ = 908.5) for *f3-cCPE-mut1*, one ([M+2H]^2+^ = 756.3) for *f1-C5C2* and one ([M+2H]^2+^ = 648.3) for *f2-C5C2*, was used as mass detection for the specific peptide. The calculation of the final concentration in each sample was performed by extrapolating it from the calibration curve using TargetLynx software (Waters®).

### Cell culture and transfection procedures

Immortalized mouse brain endothelial cells (bEnd.3) were purchased from ATCC (ATCC® CRL2299™). Cells were cultured in DMEM supplemented with 10% FBS, 1% penicillin/streptomycin, 1% glutamine. Cells were grown in T75 culture flasks and maintained in 5% CO2, 90% humidified atmosphere at 37 °C. The cell culture medium was replaced every two days, and cells were maintained between passages 25 and 30. For the monolayer, cells were seeded onto 150 μg/mL collagen-coated upper Transwell® membranes (12 mm Ø inserts, pore size 0.4 μm, growth area 1.12 cm^2^, Transwell 3460 Corning® Costar®) or onto 18 mm Ø glass coverslips at the density of 40 × 10^3^ cells per well. The cell culture medium was replaced twice a week. All reagents were purchased from Thermo Fisher Scientific, unless specified. Human embryonic kidney (HEK) cells line 293 expressing the large T antigen of simian virus 40 (SV40) (HEK 293T) were purchased from ICLC (HTL04001). Cells were cultured in DMEM supplemented with 10% FBS, 1% penicillin/streptomycin, 1% glutamine. For transfection, cells were plated on poly-D-lysine (10 μg/mL) coated Petri dishes 24 h prior to the experiment. HEK293T cells were seeded in T75 flask and transfected at 80%. To prepare lipoplexes, 100 μL of Lipofectamine2000 in DMEM was added with 40 μg pDNA (mCLDN5 tGFP-tagged, Origene #MG202442) and incubated for 20 min at room temperature (RT). The transfection complexes were added to cells and incubated for 6 h at 37 °C. The medium was then replaced by fresh medium, and cells were incubated for an additional 24 h prior to lysis and binding studies. The total protein amount of cell lysate was evaluated by measuring absorbance at 280 nm using a NanoDrop 2000 (Thermo Fisher Scientific). The total GFP amount was estimated through a calibration curve obtained by measuring the fluorescence intensity of the recombinant tGFP Protein (Pontellina plumata, CAT#: TP700079, Origene) at increasing concentrations with a Tecan SPARK (Tecan, Switzerland) multimode microplate reader.

### Cell lysate preparation

Among various lysis methods explored to maintain the highest GFP fluorescent signal, flash freeze-thawing in liquid nitrogen was selected as it performed similarly to RIPA lysis (see GFP emission spectra in **Figure S12**) but allowing to keep cells in phosphate-buffered saline (PBS), thus avoiding buffer exchange prior to MST analysis. The total protein concentration in the lysates was determined spectrophotometrically, and the GFP content was estimated through a fluorescence calibration curve of the recombinant tGFP (**Figure S13**). To exclude the presence of non-specific binding due to GFP, non-transfected HEK-293T cell lysates with a comparable concentration of spiked recombinant tGFP were used as control.

### Microscale thermophoresis

HEK293T cells transfected with mCLDN5-GFP were washed in PBS, detached with trypsin, and centrifuged (5 min, 200 x g). Pellets were resuspended in PBS and lysed through flash freeze-thawing in liquid nitrogen. Cell suspensions were then repeatedly passed through a 25-G needle (5 times), followed by 5 times through a 27-G needle, and finally centrifuged (10 min, 4,600 x g, 4 °C). To evaluate mCLDN5 binding, the peptides, diluted in PBS/Tween-20 (0.1%), were added in defined amounts as titration series to cell lysates containing mCLDN5-GFP. Samples were then loaded into Premium capillaries (Monolith^TM^ Series, #MO-K025, NanoTemper), and the fluorescence signal was measured with a Monolith NT.115, NanoTemper (Nano-BLUE laser, power 20-30%). Binding was characterized by the dissociation constant Kd and the analysis was performed using the MO.Affinity Analysis software (NanoTemper).

### Cell viability

bEnd.3 cells were treated with selected peptides (25 μM) for 24 h. After washing, cells were incubated for 5 min with Hoechst33342 (1 µM) for nuclear visualization, calcein-AM (1 µM) for cell viability, and propidium iodide (PI, 1 µM) for cell death quantification. Cell viability was quantified by using a Nikon Eclipse-80i upright epifluorescence microscope at 10X magnification. About 10 fields were randomly selected for each independent culture (n=2). Image analysis was performed using the Cell Profiler software (Broad Institute).

### Immunofluorescence

bEnd.3 or HEK293T monolayers, grown and treated either on glass coverslips or on plastic black 96-well culture plates (µ-Plate 96 Well Square, Ibidi, 89626), were fixed in 4% paraformaldehyde in PBS for 15 min at RT. Cells were permeabilized with 0.1% Triton X-100 for 5 min, then blocked with 2% bovine serum albumin (BSA) for 30 min at RT, and incubated with the primary antibody (rabbit polyclonal anti-CLDN5, Invitrogen, #PA599415, 1:500) in the same blocking solution for 3 h. After several PBS washes, cells were incubated for 60 min with the appropriate secondary antibodies (Molecular Probes, Thermo-Fisher Scientific) diluted 1:500 in the blocking buffer solution. Finally, cells were incubated for 5 min with Hoechst (1 μM, #3342, Sigma) for nuclear visualization and mounted with Vectashield antifade mounting medium (#H-1000-10, Vector Laboratories) on microscope slides. After CLDN5 immunostaining, images were acquired using a laser-scanning confocal microscope (SP8, Leica microsystem GmbH, Wetzlar, German) at 63X magnification. For each set of experiments, all images were acquired using identical exposure settings. Five fields and an equal number of z-stacks for each sample were considered. Maximal projection images were then analyzed on CellProfiler, employing a custom-made pipeline able to distinguish intracellular and extracellular CLDN5 based on intensity and morphology.

### TEER measurements and transport studies

Cells were grown until reaching a plateau in the TEER values of about 10-15 Ω·cm^2^. At this point, the apical fraction was replaced with a solution of 25 μM of peptides or vehicle. TEER values were monitored every hour for 72 h using a CellZscope+ instrument (NanoAnalytics, Münster, Germany).

Transport studies were performed by measuring the 4/70 kDa fluorescein isothiocyanate–dextran (FD4 - #46944, FD70, Sigma-Aldrich) permeability on cells at 37 °C for 2-72 h. Before each experiment, the Transwell® apical medium was replaced with 40 μg/mL FD4/70 and 25 µM f1-C5C2 or f3-cCPE-mut1 in cell medium, and the basolateral compartment was filled with 1.5 mL of fresh medium. At the indicated time points, 100 μL aliquots were sampled from the basolateral chamber into a 96-wells plate and immediately replaced by 100 μL of fresh cell medium to maintain the total volume. Fluorescence measurements were conducted using a Tecan SPARK (Tecan, Switzerland) multimode microplate reader using the 485 nm FITC channel for excitation and reading the fluorescence emission at 535 nm. Serial dilutions of FD4/FD70 in the range of 0-300 μg/ml in cell medium were prepared to obtain a calibration curve. Linear regression was applied to define the correlation between fluorescence intensity and FD4/FD70 mass concentration and used to determine the total mass of FD4/FD70 in the basolateral chamber. The regression coefficients obtained from the linear curve fits were generally 0.98-0.99 (n = 3 wells, from 3 independent culture preparations) and the apparent permeability (Papp) of FD4 was determined using the equation:

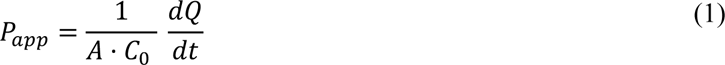

where *A* represents the surface area of the membrane in cm^2^, *C*_0_ is the initial concentration of FD4/70 in the apical compartment (μg/mL), and *dQ*/*dt* is the amount of FD4/70 in the basolateral compartment in the given time period (μg/min).

### Western blotting of TJ proteins

bEND.3 cells were grown on 6-well plates until reaching confluence and then eventually treated with 10 µM and 25 µM f1-C5C2 for 24 h before protein extraction. The day after, the medium was removed and cells were washed 3x in cold PBS. bEND.3 cells were then lysed by adding RIPA buffer (50 mM Tris-HCl pH 7.4, 150 mM NaCl, 1% Igepal, 0.1% sodium dodecyl sulfate and 0.5% sodium deoxycholate) supplemented with proteases inhibitors (complete EDTA-free protease inhibitors, Roche Diagnostic, Monza, Italy) and phosphatases inhibitors (serine/threonine phosphatase inhibitor and tyrosine phosphatase inhibitor, Sigma) and by scraping them off. Cell lysates were sonicated for 10 s (Branson SLPe; 25% amplitude) and centrifuged (4 °C, 19,000 x g, 15 min). Supernatants were collected and the total amount of proteins was calculated by the BCA Protein Assay kit (ThermoFisher Scientific) using a BSA standard curve. Samples were were denaturated for 5 min at 95 °C and separated by sodium dodecyl sulfate-polyacrylamide gel electrophoresis (SDS-PAGE; 5% and 8-12% acrylamide in the stacking and running gels, respectively). After the run, samples were transferred onto a nitrocellulose membrane (Amersham Protran, Cytiva) for 90 min at 100 V at 4 °C. Membranes were blocked with 5% milk solution in Tris-buffered saline containing 0.05% Tween-20 (0.05% TBS-T) for 1 h at RT and then incubated overnight with antibodies against: ZO-1 (rabbit polyclonal antibody, Invitrogen, #61-7300), occludin (mouse monoclonal antibody, Invitrogen, #33-1500, 1:1000), claudin 5 (rabbit polyclonal, Invitrogen, #PA599415, 1:500) and the housekeeping protein calnexin (rabbit polyclonal antibody, Enzo Life Sciences, ADI-SPA-860, 1:1000). After 3x washes with 0.05% TBS-T, membranes were incubated with HRP-conjugated secondary antibodies (Abcam, 1:10000) for 1 h at RT and developed using the ECL Prime Western Blotting System (Amersham Protran, Cytiva). The chemiluminescent signals were revealed using the iBright FL1500 Imaging System (Thermo Fisher Scientific, A44241) and band intensities were analyzed with iBright Analysis Software.

### Surface biotinylation

bEND.3 cells cultured on 6-well plates until confluence were either treated with either vehicle or *f1-C5C2* (25 µM). Twenty-four hours after the treatment, the cells were washed in cold PBS and incubated for 30 min with EZ-Link™ Sulfo–NHS–LC–biotin (1 mg/mL; ThermoFisher Scientific, 21335) in cold PBS (pH 8). Biotinylated cells were washed twice in 50 mM Tris-HCl (pH 8) and twice in cold PBS (pH 8) to eliminate unbound biotin and then lysed in RIPA buffer. A portion of the lysate supernatants was reserved for the total (input) sample, while the remaining volume was added with 100 μl of NeutrAvidin-conjugated agarose beads (ThermoFisher Scientific, 29201) and incubated for 3 h at 4 °C with constant rotation. After centrifugation, intracellular proteins were collected from the supernatants, while pellets were washed 2x in RIPA buffer and 2x in PBS before eluting the extracellularly labeled membrane proteins. Input, cytosolic and extracellular fractions were separated by SDS-PAGE, transferred to subjected to nitrocellulose membranes (Amersham Protran, Cytiva) as described above. Membranes were then incubated overnight with antibodies against CLDN5 (rabbit polyclonal, Invitrogen, #PA599415, 1:500) and the housekeeping cytosolic protein GAPDH (rabbit polyclonal, Abcam 9485, 1:5000) and the plasma membrane protein NaK1 (mouse monoclonal, Millipore, 05-369, 1:2000). Membranes were developed using the ECL Prime Western Blotting System (Amersham Protran, Cytiva) and analyzed as described above.

## Ethics Statements

In this work, no animal or human study is presented.

## Author Contributions

Authors contributions are attributed on the basis of CRediT author statement. MT: Conceptualization, Investigation, Validation, Data curation, Visualization, Writing – original draft. AB: Conceptualization, Investigation, Validation, Formal analysis, Data curation, Visualization, Writing – original draft. GA: Conceptualization, Investigation, Formal analysis, Writing – original draft, Supervision. EC: Investigation, Validation, Data curation. SV: Investigation, Validation, Data curation. VC: Investigation, Validation, Data curation. EM: Investigation, Validation, Data curation. DZC: Investigation, Validation, Data curation. AA: Investigation, Validation, Data curation. FP: Investigation, Validation, Data curation. FZ: Project administration, Funding acquisition, Writing – Review & Editing. VC: Conceptualization, Investigation, Validation, Data curation, Visualization, Writing – original draft, Supervision, Funding acquisition. LM: Conceptualization, Methodology, Writing – Review & Editing, Supervision, Project administration, Funding acquisition. FB: Conceptualization, Supervision, Writing – Review & Editing, Funding acquisition.

## Data availability

Data will be made available on request.

## ABBREVIATIONS

ABF: adaptive biasing force
ACN: acetonitrile
BBB: blood-brain barrier
b.END: brain endothelial
BSA: bovine serum albumin
cCPE: C-terminal fragment of *Clostridium Perfringens* enterotoxin
CLDN: claudin
CM: confocal microscopy
CNS: central nervous system
COM: center of mass
CV: collective variable
DMEM: Dulbecco’s Modified Eagle Medium
eABF: extended-system ABF
ECL: extracellular loop
FD4: 4 kDa fluorescein isothiocyanate–dextran
FE: free energy
FES: free energy surface
FF: force field
GBIS: Generalized-Born implicit solvent
GLUT1DS: glucose transporter 1 deficiency syndrome
HB: hydrogen bond
HEK: human embryonic kidney
K_d_: dissociation constant
KNN: K-nearest neighbor
LC: liquid chromatography
MD: molecular dynamics
MF-DOCK: multiflexible docking
ML: machine learning
MS: mass spectrometry
MST: microscale thermophoresis
P_app_: apparent permeability
PBC: periodic boundary conditions
PCA: principal component analysis
PME: particle mesh Ewald
PMF: potential of mean force
P-Pept: protein-peptide interactions
POPC: 1-palmitoyl-2-oleoyl-sn-glycero-3-phosphocholine
RMSD: root-mean square deviation
RMSE: root-mean square error
RT: room temperature
SIR: selected ion recording
SB: salt bridge
TAMD: temperature accelerated molecular dynamics
TEER: trans-endothelial electrical resistance
TJ: tight junction
TM: trans-membrane
VdW: van der Waals

## Acknowledgments

We thank Alessia Vignolo and Mattia Pini for the kind assistance at Fondazione Istituto Italiano di Tecnologia (IIT) computing center. Moreover, we are sincerely grateful to Sergio Decherchi for his assistance with the IIT computing center and useful discussions about machine learning. We are also grateful to Diego Moruzzo, Ilaria Dallorto, Rossana Ciancio, and Arta Mehilli for administrative assistance and technical help. We acknowledge the CINECA award under the ISCRA initiative (project IsB27, ID: HP10BQDIDT, granted to LM and project IsCa5, ID: HP10CQUP2X granted to AB). We also acknowledge the HPC infrastructure and the Support Team at Fondazione Istituto Italiano di Tecnologia. The project has received funding from Telethon-Italy (Grant Glut1 to FZ and FB), the European Union’s Horizon 2020 Research and Innovation Programme under Grant Agreement No. 881603 Graphene Flagship Core 3 (to FB), the Italian Ministry of Health (Ricerca Finalizzata GR-2021-12372966 to VC), and IRCCS Ospedale Policlinico San Martino, Genova, Italy (Ricerca Corrente and “5x1000” to FB and VC).

## REFERENCES

1. Greene C, Campbell M (2016) Tight junction modulation of the blood brain barrier: CNS delivery of small molecules. Tissue Barriers 4: e1138017.

2. Pardridge WM (2005) The blood-brain barrier: bottleneck in brain drug development. NeuroRx 2: 3– 14.

3. Koch H, Weber YG (2019) The glucose transporter type 1 (Glut1) syndromes. Epilepsy Behav 91: 90– 93.

4. Tang M, Park SH, De Vivo DC, et al. (2019) Therapeutic strategies for glucose transporter 1 deficiency syndrome. Ann Clin Transl Neurol 6: 1923–1932.

5. Wang D, Pascual JM, Vivo DD (2018) Glucose Transporter Type 1 Deficiency Syndrome, University of Washington, Seattle.

6. Hashimoto Y, Campbell M (2020) Tight junction modulation at the blood-brain barrier: Current and future perspectives. Biochim Biophys Acta Biomembr 1862: 183298.

7. Ogawa K, Kato N, Kawakami S (2020) Recent Strategies for Targeted Brain Drug Delivery. Chem Pharm Bull (Tokyo) 68: 567–582.

8. Bergmann S, Lawler SE, Qu Y, et al. (2018) Blood-brain-barrier organoids for investigating the permeability of CNS therapeutics. Nat Protoc 13: 2827–2843.

9. Bellavance M-A, Blanchette M, Fortin D (2008) Recent advances in blood-brain barrier disruption as a CNS delivery strategy. AAPS J 10: 166–177.

10. Goliaei A, Adhikari U, Berkowitz ML (2015) Opening of the Blood-Brain Barrier Tight Junction Due to Shock Wave Induced Bubble Collapse: A Molecular Dynamics Simulation Study. ACS Chem Neurosci 6: 1296–1301.

11. Dithmer S, Staat C, Müller C, et al. (2017) Claudin peptidomimetics modulate tissue barriers for enhanced drug delivery. Ann N Y Acad Sci 1397: 169–184.

12. Neuhaus W, Piontek A, Protze J, et al. (2018) Reversible opening of the blood-brain barrier by claudin-5-binding variants of Clostridium perfringens enterotoxin’s claudin-binding domain. Biomaterials 161: 129– 143.

13. Liao Z, Yang Z, Piontek A, et al. (2016) Specific binding of a mutated fragment of Clostridium perfringens enterotoxin to endothelial claudin-5 and its modulation of cerebral vascular permeability. Neuroscience 327: 53–63.

14. Staat C, Coisne C, Dabrowski S, et al. (2015) Mode of action of claudin peptidomimetics in the transient opening of cellular tight junction barriers. Biomaterials 54: 9–20.

15. Zwanziger D, Hackel D, Staat C, et al. (2012) A peptidomimetic tight junction modulator to improve regional analgesia. Mol Pharm 9: 1785–1794.

16. Aasen SN, Espedal H, Holte CF, et al. (2019) Improved Drug Delivery to Brain Metastases by Peptide-Mediated Permeabilization of the Blood-Brain Barrier. Mol Cancer Ther 18: 2171–2181.

17. Jones RM, Hynynen K (2019) Advances in acoustic monitoring and control of focused ultrasound-mediated increases in blood-brain barrier permeability. Br J Radiol 92: 20180601.

18. Cai Q, Li X, Xiong H, et al. (2023) Optical blood-brain-tumor barrier modulation expands therapeutic options for glioblastoma treatment. Nat Commun 14: 4934.

19. Berselli A, Benfenati F, Maragliano L, et al. (2022) Multiscale modelling of claudin-based assemblies: A magnifying glass for novel structures of biological interfaces. Comput Struct Biotechnol J 20: 5984–6010.

20. Piontek J, Krug SM, Protze J, et al. (2020) Molecular architecture and assembly of the tight junction backbone. Biochim Biophys Acta Biomembr 1862: 183279.

21. Krause G, Winkler L, Mueller SL, et al. (2008) Structure and function of claudins. Biochimica et Biophysica Acta (BBA) - Biomembranes 1778: 631–645.

22. Amasheh S, Schmidt T, Mahn M, et al. (2005) Contribution of claudin-5 to barrier properties in tight junctions of epithelial cells. Cell Tissue Res 321: 89–96.

23. Mrsny RJ, Brown GT, Gerner-Smidt K, et al. (2008) A key claudin extracellular loop domain is critical for epithelial barrier integrity. Am J Pathol 172: 905–915.

24. Kondoh M, Takahashi A, Fujii M, et al. (2006) A novel strategy for a drug delivery system using a claudin modulator. Biol Pharm Bull 29: 1783–1789.

25. Takahashi A, Kondoh M, Masuyama A, et al. (2005) Role of C-terminal regions of the C-terminal fragment of Clostridium perfringens enterotoxin in its interaction with claudin-4. J Control Release 108: 56–62.

26. Protze J, Eichner M, Piontek A, et al. (2015) Directed structural modification of Clostridium perfringens enterotoxin to enhance binding to claudin-5. Cell Mol Life Sci 72: 1417–1432.

27. Chen L, Sutharsan R, Lee JL, et al. (2022) Claudin-5 binder enhances focused ultrasound-mediated opening in an in vitro blood-brain barrier model. Theranostics 12: 1952–1970.

28. Suzuki H, Nishizawa T, Tani K, et al. (2014) Crystal structure of a claudin provides insight into the architecture of tight junctions. Science 344: 304–307.

29. Shinoda T, Shinya N, Ito K, et al. (2016) Structural basis for disruption of claudin assembly in tight junctions by an enterotoxin. Sci Rep 6: 33632.

30. Nakamura S, Irie K, Tanaka H, et al. (2019) Morphologic determinant of tight junctions revealed by claudin-3 structures. Nat Commun 10: 816.

31. Vecchio AJ, Stroud RM (2019) Claudin-9 structures reveal mechanism for toxin-induced gut barrier breakdown. Proc Natl Acad Sci U S A 116: 17817–17824.

32. Irudayanathan FJ, Wang N, Wang X, et al. (2017) Architecture of the paracellular channels formed by claudins of the blood-brain barrier tight junctions. Ann N Y Acad Sci 1405: 131–146.

33. Irudayanathan FJ, Nangia S (2020) Paracellular Gatekeeping: What Does It Take for an Ion to Pass Through a Tight Junction Pore? Langmuir 36: 6757–6764.

34. Berselli A, Alberini G, Benfenati F, et al. (2022) Computational Assessment of Different Structural Models for Claudin-5 Complexes in Blood–Brain Barrier Tight Junctions. ACS Chem Neurosci 13: 2140– 2153.

35. Weng G, Gao J, Wang Z, et al. (2020) Comprehensive Evaluation of Fourteen Docking Programs on Protein–Peptide Complexes. J Chem Theory Comput 16: 3959–3969.

36. Agrawal P, Singh H, Srivastava HK, et al. (2019) Benchmarking of different molecular docking methods for protein-peptide docking. BMC Bioinformatics 19: 426.

37. Santos KB, Guedes IA, Karl ALM, et al. (2020) Highly Flexible Ligand Docking: Benchmarking of the DockThor Program on the LEADS-PEP Protein-Peptide Data Set. J Chem Inf Model 60: 667–683.

38. Antunes DA, Devaurs D, Moll M, et al. (2018) General Prediction of Peptide-MHC Binding Modes Using Incremental Docking: A Proof of Concept. Sci Rep 8: 4327.

39. Ciemny M, Kurcinski M, Kamel K, et al. (2018) Protein–peptide docking: opportunities and challenges. Drug Discovery Today 23: 1530–1537.

40. London N, Raveh B, Schueler-Furman O (2013) Peptide docking and structure-based characterization of peptide binding: from knowledge to know-how. Curr Opin Struct Biol 23: 894–902.

41. Xu X, Zou X (2022) Predicting Protein-Peptide Complex Structures by Accounting for Peptide Flexibility and the Physicochemical Environment. J Chem Inf Model 62: 27–39.

42. Apostolopoulos V, Bojarska J, Chai T-T, et al. (2021) A Global Review on Short Peptides: Frontiers and Perspectives. Molecules 26: 430.

43. Suzuki H, Tani K, Tamura A, et al. (2015) Model for the Architecture of Claudin-Based Paracellular Ion Channels through Tight Junctions. J Mol Biol 427: 291–297.

44. Berselli A, Alberini G, Benfenati F, et al. (2023) The impact of pathogenic and artificial mutations on Claudin-5 selectivity from molecular dynamics simulations. Comput Struct Biotechnol J 21: 2640–2653.

45. Sievers F, Higgins DG (2018) Clustal Omega for making accurate alignments of many protein sequences. Protein Sci 27: 135–145.

46. Higgins DG, Thompson JD, Gibson TJ (1996) Using CLUSTAL for multiple sequence alignments, Methods in Enzymology, Academic Press, 383–402.

47. Di Marino D, Chillemi G, De Rubeis S, et al. (2015) MD and Docking Studies Reveal That the Functional Switch of CYFIP1 is Mediated by a Butterfly-like Motion. J Chem Theory Comput 11: 3401–3410.

48. D’Annessa I, Di Leva FS, La Teana A, et al. (2020) Bioinformatics and Biosimulations as Toolbox for Peptides and Peptidomimetics Design: Where Are We? Front Mol Biosci 7: 66.

49. Di Leva FS, Tomassi S, Di Maro S, et al. (2018) From a Helix to a Small Cycle: Metadynamics-Inspired αvβ6 Integrin Selective Ligands. Angew Chem Int Ed Engl 57: 14645–14649.

50. Maragliano L, Vanden-Eijnden E (2006) A temperature accelerated method for sampling free energy and determining reaction pathways in rare events simulations. Chem Phys Lett 426: 168–175.

51. Stoltz G, Vanden-Eijnden E (2018) Longtime convergence of the temperature-accelerated molecular dynamics method. Nonlinearity 31: 3748.

52. Lesage A, Lelièvre T, Stoltz G, et al. (2017) Smoothed Biasing Forces Yield Unbiased Free Energies with the Extended-System Adaptive Biasing Force Method. J Phys Chem B 121: 3676–3685.

53. Asmari M, Ratih R, Alhazmi HA, et al. (2018) Thermophoresis for characterizing biomolecular interaction. Methods 146: 107–119.

54. Jerabek-Willemsen M, André T, Wanner R, et al. (2014) MicroScale Thermophoresis: Interaction analysis and beyond. J Mol Struct 1077: 101–113.

55. Ahuja D, Sáenz-Robles MT, Pipas JM (2005) SV40 large T antigen targets multiple cellular pathways to elicit cellular transformation. Oncogene 24: 7729–7745.

56. Castagnola V, Deleye L, Podestà A, et al. (2023) Interactions of Graphene Oxide and Few-Layer Graphene with the Blood–Brain Barrier. Nano Lett 23: 2981–2990.

57. Gaillard PJ, de Boer AG (2000) Relationship between permeability status of the blood-brain barrier and in vitro permeability coefficient of a drug. Eur J Pharm Sci 12: 95–102.

58. Frost TS, Jiang L, Zohar Y (2020) Pharmacokinetic Analysis of Epithelial/Endothelial Cell Barriers in Microfluidic Bilayer Devices with an Air-Liquid Interface. Micromachines (Basel) 11: 536.

59. Shashikanth N, France MM, Xiao R, et al. (2022) Tight junction channel regulation by interclaudin interference. Nat Commun 13: 3780.

60. Hartsock A, Nelson WJ (2008) Adherens and tight junctions: structure, function and connections to the actin cytoskeleton. Biochim Biophys Acta 1778: 660–669.

61. Campbell HK, Maiers JL, DeMali KA (2017) Interplay between tight junctions & adherens junctions. Exp Cell Res 358: 39–44.

62. Del Vecchio G, Tscheik C, Tenz K, et al. (2012) Sodium Caprate Transiently Opens Claudin-5-Containing Barriers at Tight Junctions of Epithelial and Endothelial Cells. Mol Pharmaceutics 9: 2523–2533.

63. Krug SM, Amasheh M, Dittmann I, et al. (2013) Sodium caprate as an enhancer of macromolecule permeation across tricellular tight junctions of intestinal cells. Biomaterials 34: 275–282.

64. Hashimoto Y, Zhou W, Hamauchi K, et al. (2018) Engineered membrane protein antigens successfully induce antibodies against extracellular regions of claudin-5. Sci Rep 8: 8383.

65. Tachibana K, Hashimoto Y, Shirakura K, et al. (2021) Safety and efficacy of an anti-claudin-5 monoclonal antibody to increase blood–brain barrier permeability for drug delivery to the brain in a non-human primate. Journal of Controlled Release 336: 105–111.

66. Mehta RI, Carpenter JS, Mehta RI, et al. (2023) Ultrasound-mediated blood–brain barrier opening uncovers an intracerebral perivenous fluid network in persons with Alzheimer’s disease. Fluids and Barriers of the CNS 20: 46.

67. Cavaco M, Valle J, Flores I, et al. (2021) Estimating peptide half-life in serum from tunable, sequence-related physicochemical properties. Clinical and Translational Science 14: 1349–1358.

68. Dabrowski S, Staat C, Zwanziger D, et al. (2015) Redox-Sensitive Structure and Function of the First Extracellular Loop of the Cell–Cell Contact Protein Claudin-1: Lessons from Molecular Structure to Animals. Antioxid Redox Signal 22: 1–14.

69. Saitoh Y, Suzuki H, Tani K, et al. (2015) Tight junctions. Structural insight into tight junction disassembly by Clostridium perfringens enterotoxin. Science 347: 775–778.

70. Stamatovic SM, Keep RF, Wang MM, et al. (2009) Caveolae-mediated Internalization of Occludin and Claudin-5 during CCL2-induced Tight Junction Remodeling in Brain Endothelial Cells. J Biol Chem 284: 19053–19066.

71. Stamatovic SM, Keep RF, Andjelkovic AV (2011) Tracing the endocytosis of claudin-5 in brain endothelial cells. Methods Mol Biol 762: 303–320.

72. Zwanziger D, Staat C, Andjelkovic AV, et al. (2012) Claudin-derived peptides are internalized via specific endocytosis pathways. Ann N Y Acad Sci 1257: 29–37.

73. Beeman N, Webb PG, Baumgartner HK (2012) Occludin is required for apoptosis when claudin– claudin interactions are disrupted. Cell Death Dis 3: e273–e273.

74. Schlingmann B, Overgaard CE, Molina SA, et al. (2016) Regulation of claudin/zonula occludens-1 complexes by hetero-claudin interactions. Nat Commun 7: 12276.

75. Campbell M, Hanrahan F, Gobbo OL, et al. (2012) Targeted suppression of claudin-5 decreases cerebral oedema and improves cognitive outcome following traumatic brain injury. Nat Commun 3: 849.

76. Nitta T, Hata M, Gotoh S, et al. (2003) Size-selective loosening of the blood-brain barrier in claudin-5-deficient mice. J Cell Biol 161: 653–660.

77. Campbell M, Kiang A-S, Kenna PF, et al. (2008) RNAi-mediated reversible opening of the blood-brain barrier. J Gene Med 10: 930–947.

78. Berselli A, Alberini G, Benfenati F, et al. (2022) Computational study of ion permeation through claudin-4 paracellular channels. Ann NY Acad Sci 1516: 162–174.

79. Alberini G, Benfenati F, Maragliano L (2018) Molecular Dynamics Simulations of Ion Selectivity in a Claudin-15 Paracellular Channel. J Phys Chem B 122: 10783–10792.

80. Alberini G, Benfenati F, Maragliano L (2017) A refined model of claudin-15 tight junction paracellular architecture by molecular dynamics simulations. PLOS ONE 12: e0184190.

81. Feller SE, Zhang Y, Pastor RW, et al. (1995) Constant pressure molecular dynamics simulation: The Langevin piston method. J Chem Phys 103: 4613–4621.

82. Martyna GJ, Tobias DJ, Klein ML (1994) Constant pressure molecular dynamics algorithms. J Chem Phys 101: 4177–4189.

83. Huang J, Rauscher S, Nawrocki G, et al. (2017) CHARMM36m: an improved force field for folded and intrinsically disordered proteins. Nat Methods 14: 71–73.

84. Phillips JC, Hardy DJ, Maia JDC, et al. (2020) Scalable molecular dynamics on CPU and GPU architectures with NAMD. J Chem Phys 153: 044130.

85. Kuroda Y, Suenaga A, Sato Y, et al. (2016) All-atom molecular dynamics analysis of multi-peptide systems reproduces peptide solubility in line with experimental observations. Sci Rep 6: 19479.

86. Klauda JB, Venable RM, Freites JA, et al. (2010) Update of the CHARMM all-atom additive force field for lipids: validation on six lipid types. J Phys Chem B 114: 7830–7843.

87. Humphrey W, Dalke A, Schulten K (1996) VMD: visual molecular dynamics. J Mol Graph 14: 33– 38, 27–8.

88. Lapelosa M (2018) Conformational dynamics and free energy of BHRF1 binding to Bim BH3. Biophys Chem 232: 22–28.

89. Fiorin G, Pastore A, Carloni P, et al. (2006) Using Metadynamics to Understand the Mechanism of Calmodulin/Target Recognition at Atomic Detail. Biophys J 91: 2768–2777.

90. Wingbermühle S, Schäfer LV (2020) Capturing the Flexibility of a Protein–Ligand Complex: Binding Free Energies from Different Enhanced Sampling Techniques. J Chem Theory Comput 16: 4615–4630.

91. Lamothe G, Malliavin TE (2018) re-TAMD: exploring interactions between H3 peptide and YEATS domain using enhanced sampling. BMC Struct Biol 18: 4.

92. Abrams C, Bussi G (2014) Enhanced Sampling in Molecular Dynamics Using Metadynamics, Replica-Exchange, and Temperature-Acceleration. Entropy 16: 163–199.

93. Torrie GM, Valleau JP (1977) Nonphysical sampling distributions in Monte Carlo free-energy estimation: Umbrella sampling. J Comput phys 23: 187–199.

94. Fu H, Shao X, Chipot C, et al. (2016) Extended Adaptive Biasing Force Algorithm. An On-the-Fly Implementation for Accurate Free-Energy Calculations. J Chem Theory Comput 12: 3506–3513.

95. Comer J, Gumbart JC, Hénin J, et al. (2015) The Adaptive Biasing Force Method: Everything You Always Wanted To Know but Were Afraid To Ask. J Phys Chem B 119: 1129–1151.

96. Limongelli V, Bonomi M, Parrinello M (2013) Funnel metadynamics as accurate binding free-energy method. Proc Natl Acad Sci U S A 110: 6358–6363.

97. Lapelosa M (2017) Free Energy of Binding and Mechanism of Interaction for the MEEVD-TPR2A Peptide–Protein Complex. J Chem Theory Comput 13: 4514–4523.

98. Woo H-J, Roux B (2005) Calculation of absolute protein–ligand binding free energy from computer simulations. Proc Natl Acad Sci U S A 102: 6825–6830.

99. Deng Y, Roux B (2009) Computations of Standard Binding Free Energies with Molecular Dynamics Simulations. J Phys Chem B 113: 2234–2246.

100. Blazhynska M, Goulard Coderc de Lacam E, Chen H, et al. (2022) Hazardous Shortcuts in Standard Binding Free Energy Calculations. J Phys Chem Lett 13: 6250–6258.

101. Limongelli V (2020) Ligand binding free energy and kinetics calculation in 2020. WIREs Comput Mol Sci 10.

102. Maragliano L, Vanden-Eijnden E (2008) Single-sweep methods for free energy calculations. J Chem Phys 128: 184110.

103. Maragliano L, Cottone G, Ciccotti G, et al. (2010) Mapping the Network of Pathways of CO Diffusion in Myoglobin. J Am Chem Soc 132: 1010–1017.

104. Abrams CF, Vanden-Eijnden E (2012) On-the-fly free energy parameterization via temperature accelerated molecular dynamics. Chem Phys Lett 547: 114–119.

105. Alberini G, Alexis Paz S, Corradi B, et al. (2023) Molecular Dynamics Simulations of Ion Permeation in Human Voltage-Gated Sodium Channels. J Chem Theory Comput 19: 2953–2972.

106. Fiorin G, Klein ML, Hénin J (2013) Using collective variables to drive molecular dynamics simulations. Molecular Physics 111: 3345–3362.

107. Chen H, Fu H, Shao X, et al. (2018) ELF: An Extended-Lagrangian Free Energy Calculation Module for Multiple Molecular Dynamics Engines. J Chem Inf Model 58: 1315–1318.

