## Supplementary Information for "A claudin5-binding peptide enhances the permeability of the blood-brain-barrier"

### Supplementary Results

#### S1. Assessment of peptide solubility based on MD simulations and KNN classification model

To generate the training set, we selected 16 peptides, 8 soluble and 8 insoluble, each with a length comprised between 11 and 16 amino acids, from the Antimicrobial Peptide Database (https://aps.unmc.edu) [1] and subjected them to the same simulation and analysis procedure. For each peptide, we assembled a 3 × 3 × 3 grid of equally-spaced structures in explicit solvent and simulated the system for 100 ns. The values of the descriptors calculated along the simulations are reported in **Table S2** and shown in **Figure S4A.** The data points are neatly separated into two groups, with insoluble peptides characterized by large size and low number of aggregates, and low percentage of water contacts, while soluble peptides present small size and high number of aggregates and high percentage of water contacts. In **Figure S4B**, we show the results of a KNN supervised classification model using the external dataset as training after the projection of the descriptors space onto a two-dimensional PC plane. The KNN model clearly divides the PC plane according to the category of the instances found in each region, soluble or insoluble peptides. Notably, the separation between the two classes is largely due to PC1 (explaining 91% of the variance), with each descriptor contributing almost equally to this axis (32.4%, 33.6%, and 34.0% for the largest aggregate, number of aggregates, and water contacts, respectively). When applied to the peptides of our test set, the procedure successfully separates soluble from insoluble species.

#### S2. Comparison between TAMD and eABF simulations

The FE curves computed from either TAMD or eABF simulations are characterized by similar profiles, providing consistent indications on the different binding affinity of the two peptides towards mCLDN5. Indeed, eABF calculations yielded an energy minimum of ~ -6 kcal/mol at a distance of ~ 10 Å for *f1-C5C2* (**Figure S7A**, magenta line). In contrast, the FE profile for *f3-cCPE-mut1* is characterized by a barrier of ~ 9 kcal/mol monotonically decreasing to 0 kcal/mol as the COM-COM distance increases (**Figure S7A**, green line). However, in this case, the FE barrier exhibits a more pronounced reduction compared to FE profile obtained from TAMD calculations (**Figure 4A**, green line). Differences in the eABF profiles are limited to a small energy barrier for *f1-C5C2* at ~14 Å and a steeper curve for *f3-cCPE-mut1*, reaching a FE less than 2 kcal/mol already at a COM-COM distance of ~ 10 Å. Finally, a significant difference was observed in the convergence of the two methods (**Figure S8**). Concerning the error computed from the estimated FE profiles (**Figure S8C, D**), TAMD yielded RMSEs below 1 kcal/mol already after 70 ns and 25 ns for *f1-C5C2* and *f3-cCPE-mut1*, respectively. The same calculations performed over the FE gradients measured from TAMD trajectories revealed a steep decrease of the RMSE after ~ 40 ns and ~ 30 ns of MD simulation for *f1-C5C2* and *f3-cCPE-mut1*, respectively, reaching in both cases values stably lower than 0.5 kcal / (mol Å) for the remaining part of the simulation. On the other hand, a gradual reduction of the RMSE was found for eABF calculations, for which the error linearly approaching values close to 0 as the cumulative simulated time increased for both mCLDN5-peptide complexes.

The comparison between the representative configurations selected from TAMD (**Figure 4**) and eABF (**Figure S7)** revealed similar poses for clusters 1, 2 and 3. Indeed, interactions reported in **Table S3** show that these structures (**Figure 4D-F** for TAMD and **Figure S7D-F** for eABF) share a similar binding pattern, involving residues K48, V55, V66, E68, S69, V70 and L71 belonging to ECL1, and P153, V154 and Y158 located on ECL2. This interface is also highly comparable to the structure provided by our standard MD runs (**Figure 4B**). On the contrary, representative structures mapping higher RMSD values significantly diverged between the two methods (**Figure 4G-I** for TAMD and **Figure S7G-I** for eABF), and they are also characterized by different binding configurations with respect to structures from clusters 1, 2 and 3 (**Table S3**).

### Supplementary Methods

#### S3. Design of cCPE and C5C2-based peptides dataset

**Fragments of cCPE *(f1-cCPE/ f2-cCPE/ f3-cCPE).*** Inspired by the previously illustrated modeling protocol [2], we prepared a homology-based all-atom structural model of the hCLDN5-cCPE complex using, as template, the experimentally solved structure of the CLDN4-cCPE system (PDB ID: 5B2G [3]). The cCPE domain was modified, by including three mutations (N218Q, Y306W and S313H) that were reported to improve the binding specificity to CLDN5 [2]. After the refinement of the hCLDN5-cCPE complex with the FG-MD server [4], we extracted three peptides from this system (named as *f1-cCPE, f2-cCPE* and *f3-cCPE,* sequences in **Table S1**), by removing all the cCPE residues that have at least an atom at a distance larger than 7.5 Å from the hCLDN5 subunit.

**Mutant fragments of cCPE (*f1-cCPE-mut1/ f1-cCPE-mut2/ f2-cCPE-mut1/ f3-cCPE-mut1*).** Four additional cCPE-based peptides mutants were designed with the purpose of enhancing solubility in biological media. These new sequences (named *f1-cCPE-mut1/f1-cCPE-mut2/f2-cCPE-mut1/f3-cCPE-mut,* see **Table S1)** were generated by introducing single point mutations in the fragments described above. The GRAVY coefficient for hydropathicity, provided by the ProtParam Tool of Expasy web server [5], was adopted for a preliminary assessment of their solubility.

**C5C2 fragment (*f1-C5C2, f2-C5C2*) and its mutants *(f1-C5C2-mut1/ f1-C5C2-mut2/ f1-C5C2-mut3/ f1-C5C2-mut4/ f1-C5C2-mut5/ f1-C5C2-mut6/ f1-C5C2-mut7/ f1-C5C2-mut8).*** The *f1-C5C2* and *f2-C5C2* peptides were extracted from the previously described sequence of the C5C2 domain [6]. The latter is a mCLDN5-binding modified version of the 30 amino acid-long C1C2 peptide originally introduced to bind CLDN1 [7], and belongs to the ECL1 domain. Consistently with the procedure described above for the mutated cCPE fragments, mutations were induced in *f1-C5C2* in order to enhance its solubility in water. Since *f1-C5C2* was extracted from the CLDN5 murine sequence (UNIPROT: O54942) and contains the previously proposed S74N mutation [6], we also introduced in our dataset the murine WT variant (*f1-C5C2-mut6*) and the respective fragments derived from the human CLDN5 sequence (UNIPROT: O00501). The latter peptides were labelled as *f1-C5C2-mut7* and *f1-C5C2-mut8* for the WT and the S74N forms, respectively (**Table S1**). By contrast, *f2-C5C2* differs from *f1-C5C2* in that it lacks the two initial residues of the sequence (glutamate and serine). Given that this difference increases its hydrophobicity, we hypothesized that *f2-C5C2* could have a lower solubility in water compared to *f1-C5C2*. For this reason, at this stage, we did not insert further mutations into *f2-C5C2.*

#### S4. Modeling CLDN5 multimeric structures

To build CLDN5 multi-pore configurations, we modeled the hCLDN5 monomer by homology modeling using SWISS-MODEL [8] and the crystallographic mCLDN15 structure (PDB ID: 4P79 [9]) as a template, after refinement for the multi-protein system [10]. A triple pore hCLDN5 WT assembly was generated starting from the multi-pore configuration proposed for mCLDN15 [11]. A different configuration was obtained based on the CLDN-CLDN dimeric *cis*-interface introduced in [12], and named *back-to-back* [13]. As described in our previous works [14,15], this configuration was reproduced through protein-protein docking simulations of two CLDN5 monomers with MEMDOCK [16]. The resulting dimer was then used to generate the triple-pore arrangement with VMD 1.9.3 [17]. Finally, these structures were relaxed using GalaxyRefineComplex [18], to remove potential steric clashes among the various hCLDN5 subunits.

#### S5. Peptide structure modeling and equilibration

All the peptide structures were predicted *de novo* using PEP-FOLD3 [19]. For each peptide, we selected the first model provided by the software. This was used as starting configuration in equilibration runs of standard MD simulation performed with the Generalized Born Implicit Solvent (GBIS) method [20,21]. In GBIS, periodic boundary conditions (PBCs) are excluded and the cutoff for the van der Waals (VdW) interactions is set to 14 Å. MD simulations were performed with a 0.3 M ion concentration and a 2 fs time step. After an initial energy minimization, a trajectory of 5 ns was produced for each peptide at a temperature T = 303.15 K. The final configurations were used as the initial structure for all-atom MD simulations in explicit TIP3P [22] solvent. Each oligomer was embedded in a cubic box of ~ 45 × 45 × 45 $Å^3$, solvated with water and a physiological concentration of 0.15 M KCl. Topology and coordinates of the systems, along with the input files for the MD simulation, were generated with CHARMM-GUI [23]. After an initial equilibration performed with positional restraints on the peptide backbone, 100 ns of MD simulation were produced in the absence of constraints. These trajectories were produced in the NPT ensemble at T=303.15 K and P=1 bar, maintained by a Langevin thermostat and Nosé-Hoover Langevin barostat [24,25]. PBCs were adopted to replicate the system and remove box surface effects. Long-range electrostatic interactions were computed using the Particle Mesh Ewald (PME) algorithm [26]. Electrostatic and VdW interactions were calculated with a cutoff of 12 Å as prescribed by the CHARMM force field. A switching function was applied, starting to take effect at 10 Å to obtain a smooth decay as indicated in Ref. [27]. Chemical bonds involving hydrogen atoms and protein heavy atoms were constrained with SHAKE [28], while those of water molecules were kept fixed with SETTLE [29]. NAMD 3.0 [30] with CHARMM36m force field [31] was used to perform the simulation.

#### S6. Computational assessment of peptide water solubility

To estimate the solubility of peptides in a solvent that mimics biological media, we performed all-atom MD simulations of multiple copies of each peptide and evaluated their attitude to forming aggregates, following the previously proposed protocol [32]. The configurations from the MD simulations of each single peptide at 50, 75, and 100 ns were used to assemble a 3 × 3 × 3 grid of equally-spaced structures. Peptides were positioned at a distance of 25 Å from each other to minimize initial interactions and the N/C-termini of each peptide were neutralized with an amide and ester group, respectively. The system was solvated with explicit TIP3P [22] water molecules and neutralized with a heterogeneous 0.09 M KCl and 0.03 M $CaCl_2$ ionic bath that reproduced the ionic strength of the cell culture medium.

After an initial equilibration, each system was simulated for 100 ns. MD simulations were performed with the same set-up previously described for the single peptides, at 303.15 K. To evaluate solubility, the MD trajectories were analyzed using an *in-house* Tcl script to compute the number and dimension of the aggregates formed over time. Specifically, we assigned a peptide to an aggregate if the distance between the respective centers of mass (COM) was below the sum of the gyration radius of that peptide and the aggregate plus a threshold of 2.8 Å. Additionally, we also computed the number of contacts formed between peptides and water molecules at the end of each simulation and reported the result in percent of the contact number for the initial configuration of the system. The classification of peptides into either *soluble* or *insoluble* categories based on the reported descriptors would be ambiguous in the absence of a reference. For this reason, we replicated the same protocol for an external dataset of 16 sequences, selected from the Antimicrobial Peptide Database (https://aps.unmc.edu [1]) according to their hydrophilicity. Specifically, we identified 8 hydrophilic (hydrophilic residues > 70%) and 8 hydrophobic (hydrophilic residues < 20 %) peptides, that were used to train a supervised learning classification model based on the K-Nearest Neighbor (KNN) algorithm [20] (k=3, uniform weight function). The method was implemented with the python *scikit-learn* library [21]. The average dimension of the largest aggregate, the number of aggregates computed in last 10 ns of each trajectory, and the percentage of water contacts were used as independent variables to train the model. The dataset features were standardized onto a unit scale (mean = 0 and standard deviation = 1) and principal component analysis (PCA) was performed to create a set of uncorrelated variables in the new reference system. This model was used to predict the water solubility of peptides studied in this work. Only water-soluble oligomers were considered for subsequent steps. The workflow described in this paragraph is summarized in **Figure S2**.

#### S7. MD simulations of the mCLDN5 monomer

In the simulations, we used equilibrated structures of the mCLDN5 monomer extracted from the MD runs performed by us and described in [14]. Briefly, we generated two alternative mCLDN5 configurations by homology modeling with SWISS-MODEL [8] using the mCLDN15 (PDB ID: 4P79 [9]) and hCLDN4 (PDB ID: 5B2G) crystal structures as templates. Each model was refined with ModRefiner [33] and embedded in a pure 1-palmitoyl-2-oleoyl-sn-glycero-3-phosphocholine (POPC) bilayer, solvated with explicit TIP3P [22] water molecules and a 0.15 M KCl ionic bath. The disulfide bond between residues C54 and C64 found in the ECL1 domain was preserved. The full-hydrogen CHARMM PDB file was generated with CHARMM-GUI [23,34]. After 12-ns equilibration, a trajectory of 110 ns of production was performed in the NPT ensemble at the same temperature (310 K) and pressure (1 bar) of equilibration steps maintained by a Langevin thermostat and Nosé-Hoover Langevin piston pressure control [24,25]. Rectangular PBCs were used to replicate the system and remove box surface effects. Long-range electrostatic interactions were computed using PME [26]. Electrostatic and VdW interactions were calculated with a cutoff of 12 Å as prescribed by the CHARMM force field. A switching function was applied, starting to take effect at 10 Å to obtain a smooth decay [27]. Hydrogen atoms not involved in covalent bonds with water were restrained with SHAKE [28], while those of water molecules were kept fixed with SETTLE [29]. The NAMD 3.0 program [30] with the CHARMM36m force field [35] was adopted in these MD simulations.

#### S8. Preliminary prediction of the mCLDN5-peptide binding configuration

Two peptides (*f1-C5C2* and *f3-cCPE-mut1*), whose water solubility was predicted in the previous steps, were tested for their affinity for mCLDN5. To provide an initial guess of the binding pose of the peptide with the extracellular domain of the protein, we used AutoDock Vina [36]. This software allows the confinement of the conformational search in a selected region of the protein. Specifically, we imposed the binding in a rectangular region of $35 \times 40 \times 30$ Å$^3$ centered in the COM of the mCLDN5 extracellular domain. The required PDBQT format files were generated with the AutoDock tools suite [37,38], while results were visualized and converted to PDB with UCSF Chimera [39]. Since AutoDock Vina is optimized to manage a maximum of 32 torsional degrees of freedom for the ligand (optimal 20) [36], and due to the greater number of torsional angles of our peptides, we identified the binding pose using a stepwise procedure. First, each oligomer was docked by allowing the movement of the side chains of the first 4-5 amino acids of the sequence according to the number of rotamers, while permitting only rigid translations for the other residues. After the first iteration, the resulting peptide configuration was used for a following run performed by fixing the bonds that were sampled in the first step, and by releasing the successive. The procedure was reiterated until each torsional degree of freedom was sampled. The best pose from each cycle was selected according the to the root-mean square deviation (RMSD) value calculated with respect to the configuration resulting from the previous step. The final structure for each mCLDN5-peptide complex was further refined with FlexPepDock [40,41]. Because of this limitation of the Vina scoring function in handling the peptide flexibility [42,43], molecular docking calculations were performed only to obtain a starting configuration of each complex. Structures generated in this way have been subsequently investigated by MD simulations.

#### S9. Standard MD simulations and analysis of the mCLDN5-peptides complexes

Selected mCLDN5-peptide configurations, provided by the preliminary docking protocol, were embedded in a POPC bilayer and solvated with explicit TIP3P [22] water molecules and a 0.15 M KCl ionic bath. Analogously to hCLDN5, the disulfide bridge between the mCLDN5 residues C54 and C64 found in the ECL1 domain was preserved. The CHARMM PDB file of each protein-peptide system was generated using the CHARMM-GUI PDB manipulator [23,34], while topology files were built with the *psfgen* tool of VMD 1.9.3 [17]. After a short 50-ps-long minimization and 30 ns of equilibration with a progressive release of positional restraints on the heavy atoms of the system, 200 ns of MD simulations were performed for both the mCLDN5-*f1-C5C2* and the mCLDN5-*f3-cCPE-mut1* systems. The trajectories were generated in the NPT ensemble at the same temperature (310 K) and pressure (1 bar) of equilibration steps, maintained by a Langevin thermostat and the Nosé-Hoover Langevin piston pressure control [24,25]. The oscillation period of the piston was set to 50 fs, the damping time scale to 25 fs and the damping coefficient of the Langevin thermostat to 1 ps $^{-1}$. Rectangular PBCs were used to replicate the system and remove box surface effects. Long-range electrostatic interactions were computed using PME [26]. Electrostatic and VdW interactions were calculated with a cutoff of 12 Å and the application of a switching function for smooth decay starting at 10 Å [27]. Hydrogen atoms not involved in covalent bonds with water were restrained with SHAKE [8], while those of water molecules were kept fixed with SETTLE [29], thus enabling the use of a time step of 2 fs. The NAMD 3.0 program [30] with the CHARMM36m force field [35] were used together with the TIP3P [22] water model and the associated ionic parameters with the NBFIX corrections [44–46]. During production, the protein was restrained from lateral movement, and the peptide rearrangement was confined in a cylindrical region of radius 30Å and height 80 Å, centered at (-2.0 Å, 0.0 Å, 30.0 Å) to prevent lateral diffusion, as shown in **Figure S5**. The two MD trajectories were analyzed with VMD 1.9.3 [17] by computing structural descriptors of the mCLDN5-peptide interactions, including the presence and persistence time of hydrogen bonds (HB), salt bridges (SB) and the backbone root-mean square deviation (RMSD) of both the peptide and the mCLDN5 ECL domain. The HB were defined with a cutoff distance between the heavy heteroatoms (nitrogen, oxygen) covalently bound to a hydrogen atom of 3.3 Å, and an angle defined by the heavy atoms and the central hydrogen atom comprised between 130° and 230°. SB were computed between the charged atoms of amino acids with the opposite charge at a distance < 4.0 Å and sharing a HB. Only HB with a persistence time greater than 20% of the whole trajectory were considered a significative hallmark of the mCLDN5-peptide interactions. In addition, the number of contacts between each simulated peptide and the mCLDN5 extracellular domain was calculated with the Colvars module [47] implemented in NAMD. Specifically, the number of contacts is calculated over two groups of atoms (*group1* and *group2*), which are defined by the heavy atoms of the peptide and the mCLDN5 extracellular domain, respectively, using the following function:

$$C(group1, group2) = \sum_{i \in group1} \sum_{j \in group2} \frac{1 - \left(|\boldsymbol{r}_i - \boldsymbol{r}_j|/d_0\right)^n}{1 - \left(|\boldsymbol{r}_i - \boldsymbol{r}_j|/d_0\right)^m} \quad \text{(S1)}$$

where $\boldsymbol{r}_i$ and $\boldsymbol{r}_j$ are the cartesian coordinates of the atoms $i$ and $j$, respectively. Moreover, $n$ and $m$ are integers that control the long-range decay and stiffness of the function, and they have values of 6 and 12, respectively, while $d_0$ is a cutoff distance (here set at 4 Å).

#### S10. Enhanced MD simulations

**Temperature-accelerated MD simulations.** Temperature-accelerated MD (TAMD) [48] allows sampling of infrequent events by accelerating the dynamics of a set of auxiliary variables tethered to functions of the system's Cartesian coordinates (usually called collective variables, CVs). Let us indicate with $\boldsymbol{x} \in \mathbb{R}^{3N}$ the coordinates of the system, with $\boldsymbol{\theta} = \theta_1, \dots, \theta_M$ the CVs and with $\boldsymbol{z} = z_1, \dots, z_M$ the auxiliary variables, with $M \ll N$. The extended system $(\boldsymbol{x}, \boldsymbol{z})$ is subjected to the following potential:

$$U_k(\boldsymbol{x}, \boldsymbol{z}) = V(\boldsymbol{x}) + \frac{1}{2} k \sum_{\alpha}^{M} (\theta_\alpha(\boldsymbol{x}) - z_\alpha)^2 \quad \text{(S2)}$$

where $V(\boldsymbol{x})$ is the MD force field and $k > 0$ is a constant. As a consequence, the evolution of the extended system can be described by a set of equations such as for example:

$$\begin{cases} m\ddot{\boldsymbol{x}} = -\nabla V(\boldsymbol{x}) - k \sum_{\alpha}^{M} (\theta_\alpha(\boldsymbol{x}) - z_\alpha) \nabla \theta_\alpha(\boldsymbol{x}) + \textit{thermostat at temperature } T \\ \bar{\gamma}\dot{\boldsymbol{z}} = k(\theta(\boldsymbol{x}) - z) + \sqrt{2k_B \bar{T} \bar{\gamma}}\ \boldsymbol{\eta}(t) \end{cases} \quad \text{(S3)}$$

In these equations, $m$ is the mass matrix, $\boldsymbol{\eta}(t)$ is a Gaussian process with mean zero and covariance that $\langle \boldsymbol{\eta}_\alpha(t)\, \boldsymbol{\eta}_{\alpha'}(s) \rangle = \delta_{\alpha\alpha'}\delta(t-s)$, $\bar{\gamma}$ is an artificial friction coefficient and $\bar{T}$ is an artificial temperature. The main point of TAMD is that by adjusting $\bar{\gamma}$ so that the $\boldsymbol{z}(t)$ evolves slower than the $\boldsymbol{x}(t)$, and by tuning the parameter $k$ so that $\boldsymbol{z}(t) \sim \boldsymbol{\theta}(\boldsymbol{x}(t))$, one can obtain a trajectory $\boldsymbol{z}(t)$ that moves at the artificial temperature $\bar{T}$ on the free energy surface (FES) calculated at the physical $T$ [48,49]. Indeed, under such assumptions, the force that drives each auxiliary variable $z_\alpha$ in eq. (S2) satisfies:

$$k(\theta_\alpha(\boldsymbol{x}) - z_\alpha) \approx -\frac{\partial G(\boldsymbol{z})}{\partial z_\alpha} \tag{S4}$$

where $G(\boldsymbol{z})$is the FE defined at the physical temperature $T$. Therefore, by choosing $\bar{T} > T$, the system will rapidly visit regions where FE is relatively low, overcoming barriers that would take a long time to be crossed at the physical $T$. In our case, we chose the distance between the peptide and the mCLDN5 ECL COMs as CV, and mapped in a range 8 to 30 Å. The full CV range was then split into two independent windows, from 8 to 20 Å and from 20 to 30 Å, respectively, and each window was simulated for 150 ns for both *f1-C5C2* and *f3-cCPE-mut1* systems. Convergence of the calculation was estimated by computing the root-mean square error (RMSE) of the FE profile or FE gradients (i.e., minus the mean forces) calculated every 10 ns of MD simulation with respect to those at 150 ns. The equilibrated mCLDN5-*f1-C5C2* and mCLDN5-*f3-cCPE-mut1* structures, already used for the standard MD simulation, were selected as starting configurations, and three independent calculations were performed for each system. As for the MD simulation parameters, the TAMD runs were carried out with the same set-up of the standard simulations previously described. The instantaneous force introduced in eq. (S4) was collected in bins of 0.1 Å. A fictitious temperature $\bar{T}$= 1000 K and a friction coefficient $\bar{\gamma}$ = 15 ps$^{-1}$ were used for to the CV's dynamics. The FE was computed by integrating the average force with the corrected z-averaged (CZAR) estimator [50]. The FE profile and the associated error were reported as the mean and the standard deviation from the three replicates, respectively.

**Extended-system adaptive biasing force.** To validate the results of the TAMD simulations, we calculated the same FE using a different enhanced sampling approach, the extended-system adaptive biasing force (eABF) method [50,51]. In eABF, the dynamics of a set of auxiliary variables restrained to the CVs is also introduced, and their sampling is achieved by adding force that counteracts the average force acting on them and by increasing their temperature [52]. eABF calculations were performed using the NAMD Colvars module [47] with the same set-up of TAMD trajectories. The CV was again the distance between the peptide and the mCLDN5 ECL COMs, and its artificial temperature was set at the same temperature of the physical system (T = 310 K). The instantaneous force was collected in bins of 0.1 Å and 1000 samples were collected before applying the biasing force. The FE was computed by integrating the average force with the CZAR estimator [50]. The FE profile and the associated error were reported as the mean and the standard deviation from three independent trajectories.

**Calculation of dissociation constant from the FE profile.** The approach to calculate the equilibrium dissociation constant, $K_d$, from MD simulations has been discussed in several works [53–59] and successfully applied to protein-ligand [57,59–63] and protein-peptide [64,65] systems. In the presence of a cylindrical restraint that confines the movement of the peptide [53,66], the $K_d$ can be estimated as follows:

$$(Kd)^{-1} = C^0(\pi R^2)\int_{r_{min}}^{r_{max}} e^{-\frac{G(r)-G(r^*)}{k_B T}}\, dr \tag{S5}$$

Where $G(r)$ is the FE calculated at the distance $r$ between the protein and the peptide, while $r^*$ is the distance at which the peptide is found in the solution bulk, completely dissociated from the protein. $C^0$ is the standard molar concentration ($\sim 1/1660$ Å$^{-3}$), $R$ is the radius of the cylindrical restraint (here 30 Å), $k_B$ is the Boltzmann's constant and $T$ the temperature (310 $K$). The integral was numerically calculated along the CV values range, from $r_{min}$ = 8 Å to $r_{max}$ = 30 Å (here, $r^* \equiv r_{max}$) adopting a bin width of 0.1 Å.

### Supplementary Figures

```
sp|O00501|CLD5_HUMAN      MGSAALEILGLVLCLVGWGGLILACGLPMWQVTAFLDHNIVTAQTTWKGLWMSCVVQSTG  60
sp|O54942|CLD5_MOUSE      MGSAALEILGLVLCLVGWVGLILACGLPMWQVTAFLDHNIVTAQTTWKGLWMSCVVQSTG  60
                          ****************** *****************************************

sp|O00501|CLD5_HUMAN      HMQCKVYDSVLALSTEVQAARALTVSAVLLAFVALFVTLAGAQCTTCVAPGPAKARVALT  120
sp|O54942|CLD5_MOUSE      HMQCKVYESVLALSAEVQAARALTVGAVLLALVALFVTLTGAQCTTCVAPGPVKARVALT  120
                          *******:******:**********.*****:*******:************.*******

sp|O00501|CLD5_HUMAN      GGVLYLFCGLLALVPLCWFANIVVREFYDPSVPVSQKYELGAALYIGWAATALLMVGGCL  180
sp|O54942|CLD5_MOUSE      GGALYAVCGLLALVPLCWFANIVVREFYDPTVPVSQKYELGAALYIGWAASALLMCGGGL  180
                          **.** .***********************:*******************:**** ** *

sp|O00501|CLD5_HUMAN      LCCGAWVCTGRPDLSFPVKYSAPRRPTATGDYDKKNYV      218
sp|O54942|CLD5_MOUSE      VCCGAWVCTGRPEFSFPVKYSAPRRPTANGDYDKKNYV      218
                          :***********::**************.*********
```

***Figure S1. Sequence alignment between human (UNIPROT: O00501) and mouse (UNIPROT: O54942) CLDN5***. *The alignment was performed with Clustal. Residues are colored according to their charge or polarity: red for apolar, green for polar, blue for acidic and magenta for basic residues.*

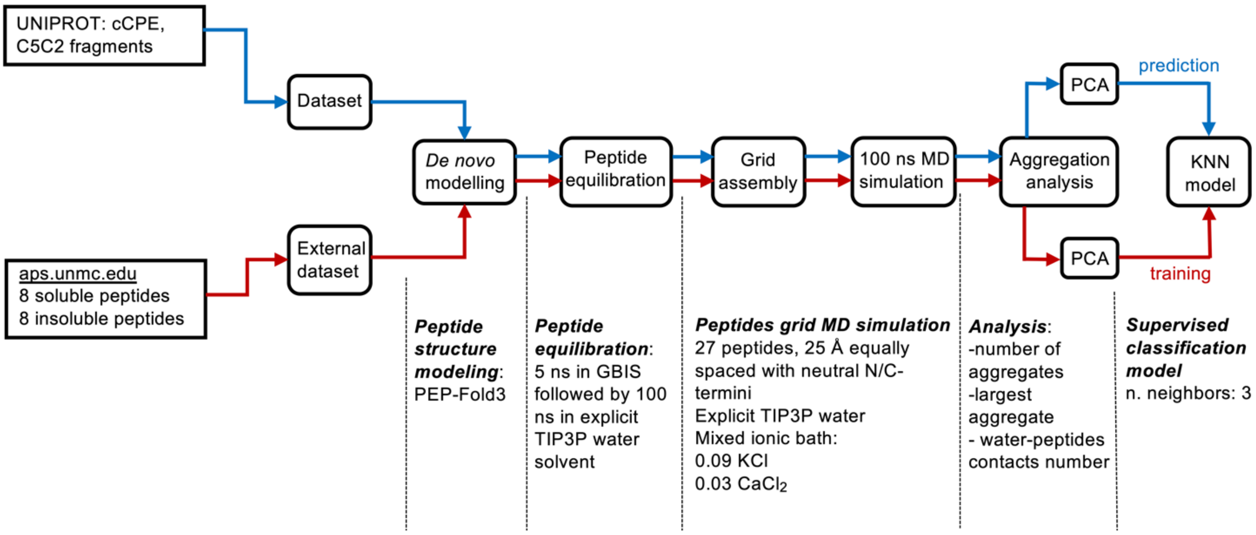


***Figure S2. Workflow of the modeling strategy to assess peptide solubility***. *The strategy is based on structural modeling, MD simulations and machine learning analysis.*

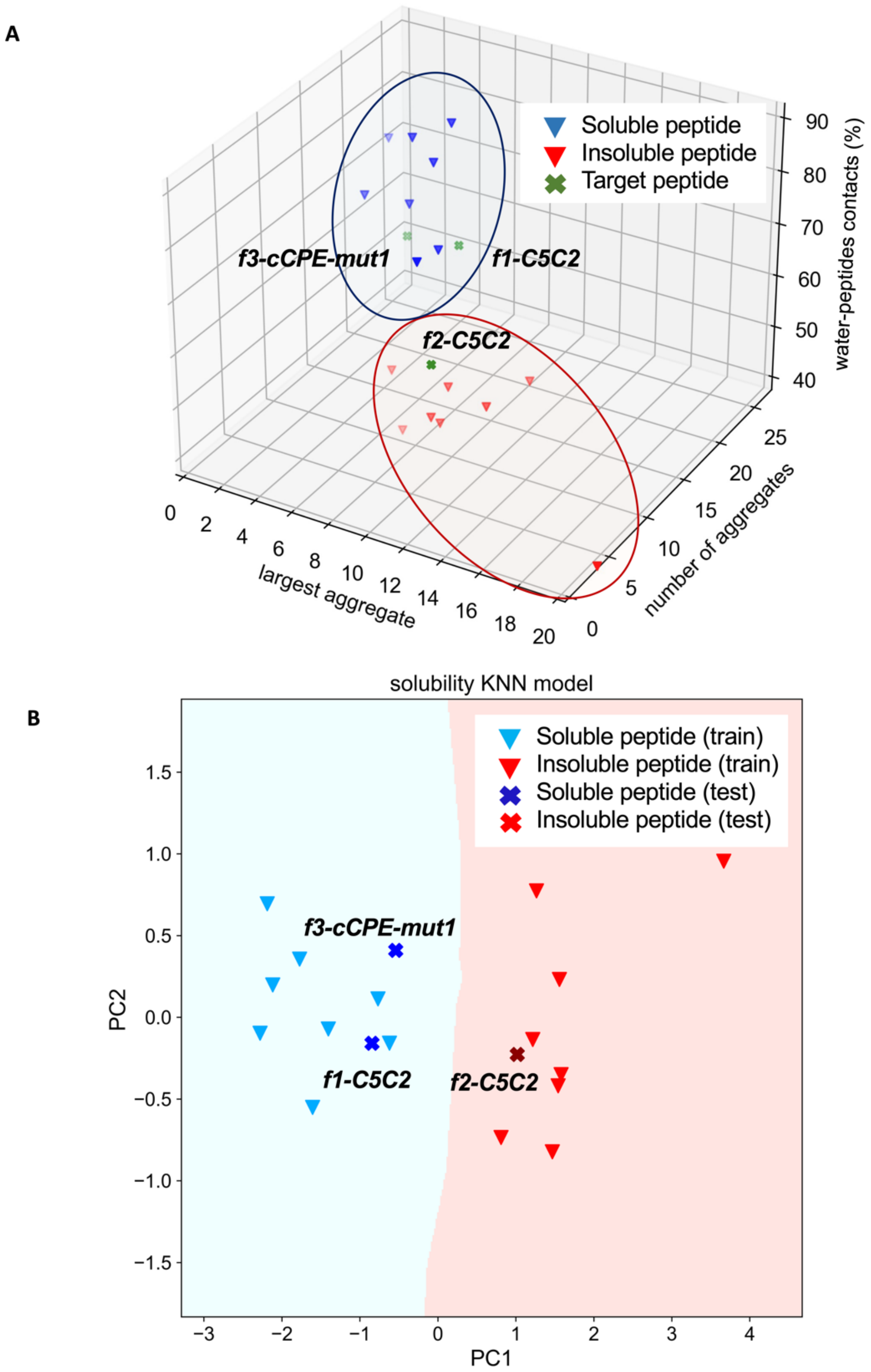


***Figure S3. Solubility classification of peptides based on the external dataset.*** ***A.*** *Three-dimensional plot of the descriptors computed during the MD simulations: the number of peptides forming the largest aggregate, the number of different aggregates and the water-peptides contacts number expressed as a percentage with respect to the starting configuration. Peptides of the external dataset are indicated as blue and red triangles for soluble and insoluble oligomers, respectively. Peptides studied in this work are shown as green crosses.* ***B.*** *Solubility prediction based on the K-nearest neighbor (KNN) model. Data dimensionality was reduced with PCA. The PC-plane is divided into two regions: peptides in the blue area are included among the soluble oligomers, while those found in the red area are included among the insoluble ones. The KNN model was trained with the external peptide dataset, while the peptides studied in this work were classified as soluble/insoluble according to their position on the PC-plane. Peptides of the external dataset are indicated as blue and red triangles for soluble and insoluble oligomers, respectively. The peptides studied in this work are shown as crosses and colored according to the position on the PC-plane.*

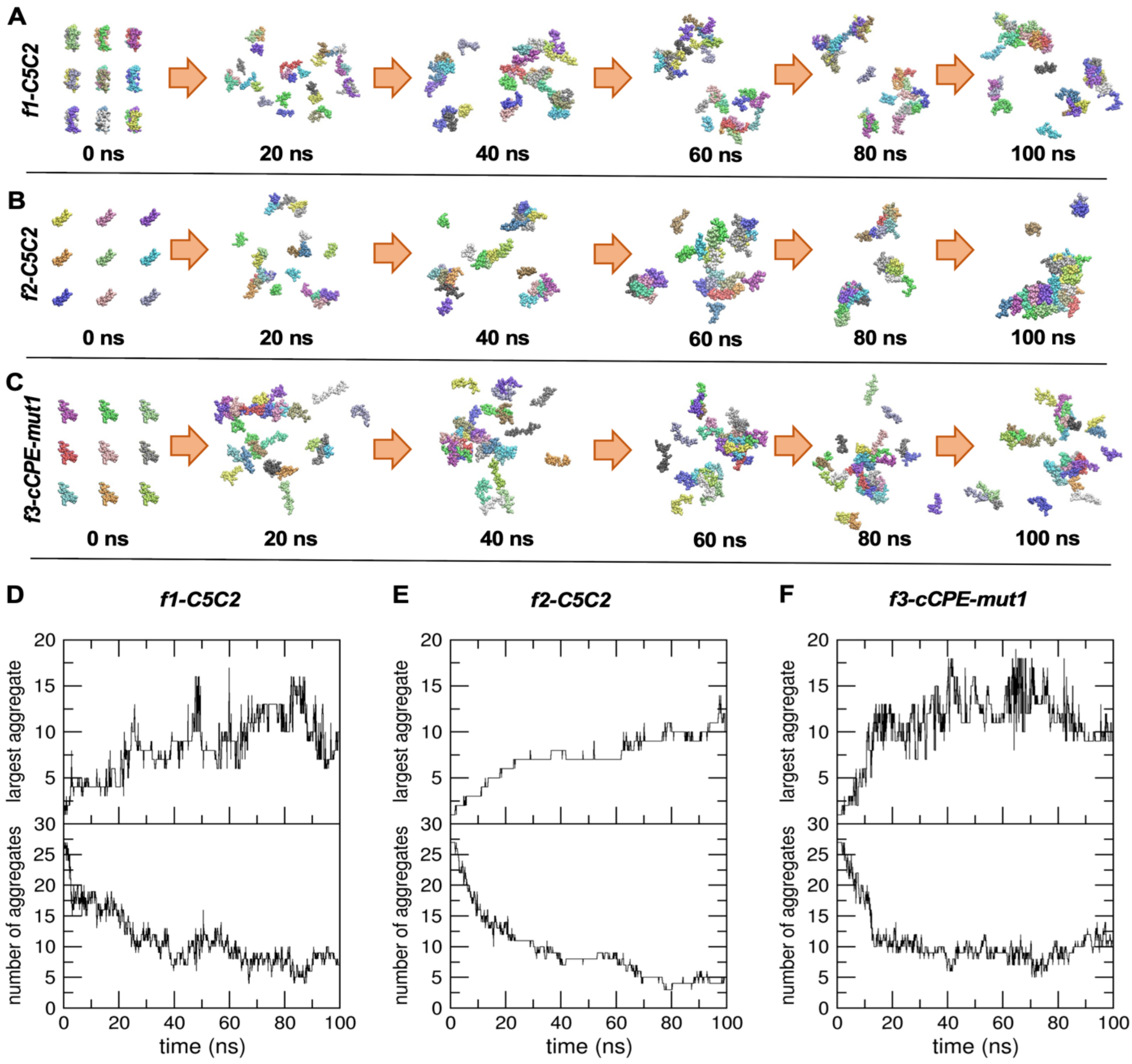


***Figure S4. MD simulations for the computational assessment of peptides water solubility.*** *Snapshots every 20 ns of the simulated trajectory are shown for the f1-C5C2 (A), f2-C5C2 (B) and f3-cCPE-mut1 (C) systems. Atoms are indicated as VdW spheres and peptides of each system are distinguished by distinct colors. Water solvent and ions are omitted for clarity. The dimension of the largest aggregate (upper panel) and the number of aggregates (lower panel) computed during the trajectories for f1-C5C2 (D), f2-C5C2 (E) and f3-cCPE-mut1 (F) are reported.*

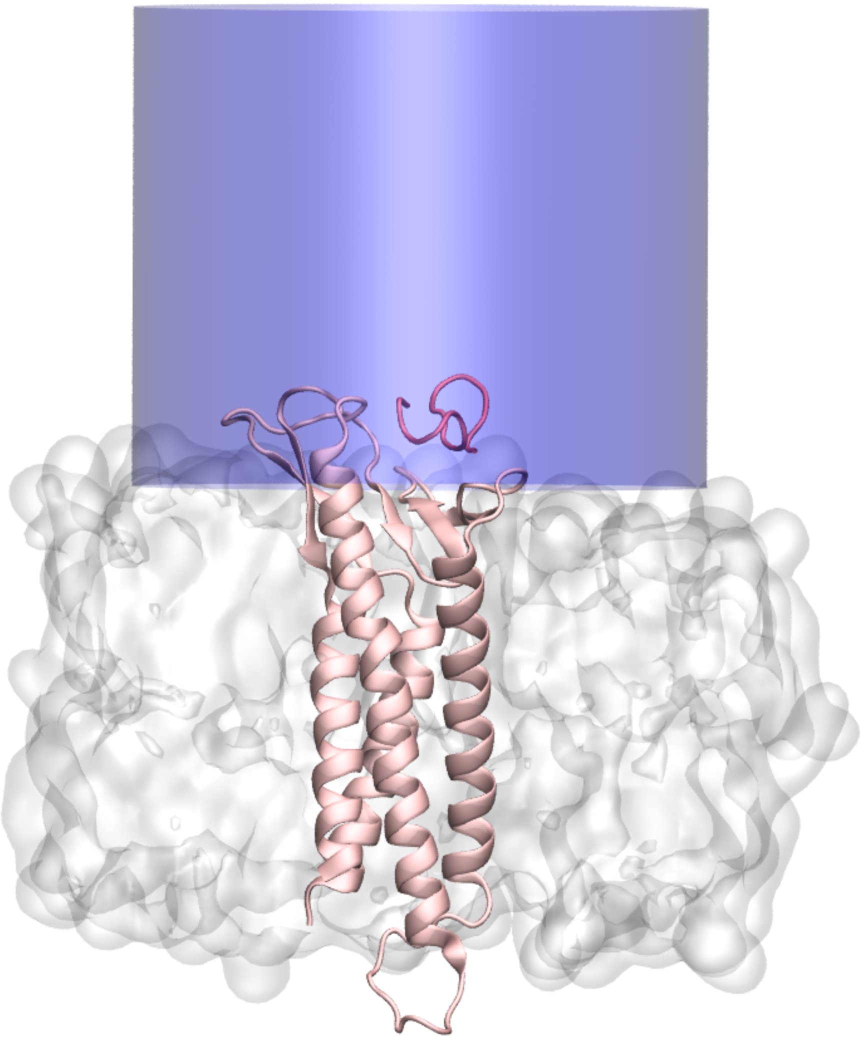

***Figure S5. Graphical representation of the cylindrical restraint used for MD simulations of protein-peptide complexes.*** *The peptide (red) and mCLDN5 (pink) embedded in the POPC bilayer (gray) are shown. The peptide movement is confined in a cylindrical volume (blue region) of radius 30Å and height 80 Å, centered at (-2.0 Å, 0.0 Å, 30.0 Å), to prevent lateral diffusion and the contact with the mCLDN5 intracellular domain.*

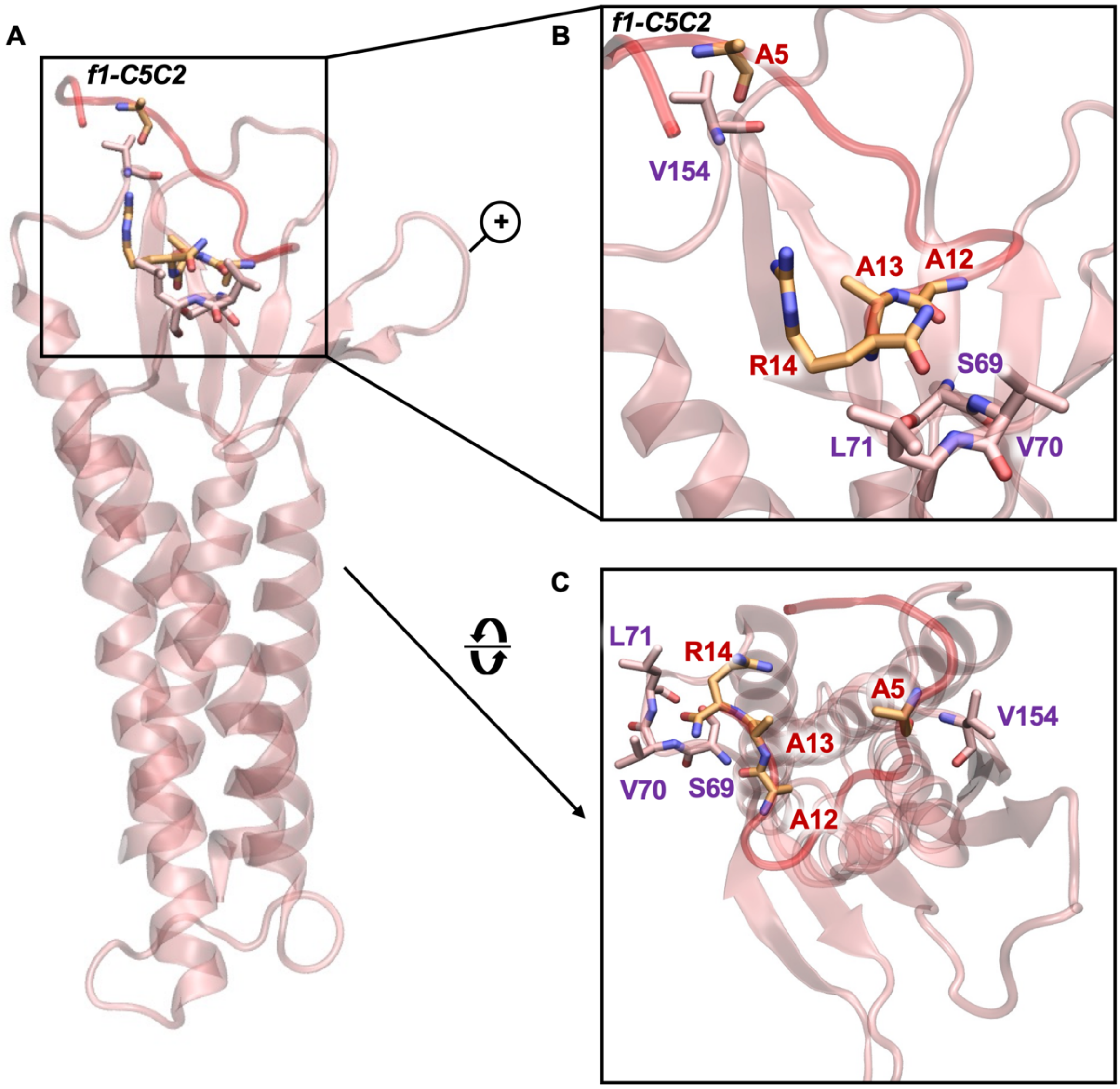


***Figure S6. Residues involved in the formation of stable HBs during MD simulations. A.*** *CLDN5-f1-C5C2 complex after 200 ns of standard MD simulations.* ***B.*** *Close-up view of the CLDN5 ECL domain. Residues involved in the formation of persistent HBs are indicated.* ***C.*** *Upper view of the CLDN5-f1-C5C2 complex. Amino acids belonging to CLDN5 are shown as pink sticks, those belonging to f1-C5C2 are colored in orange. Nitrogen and oxygen atoms are colored in blue and red, respectively.*

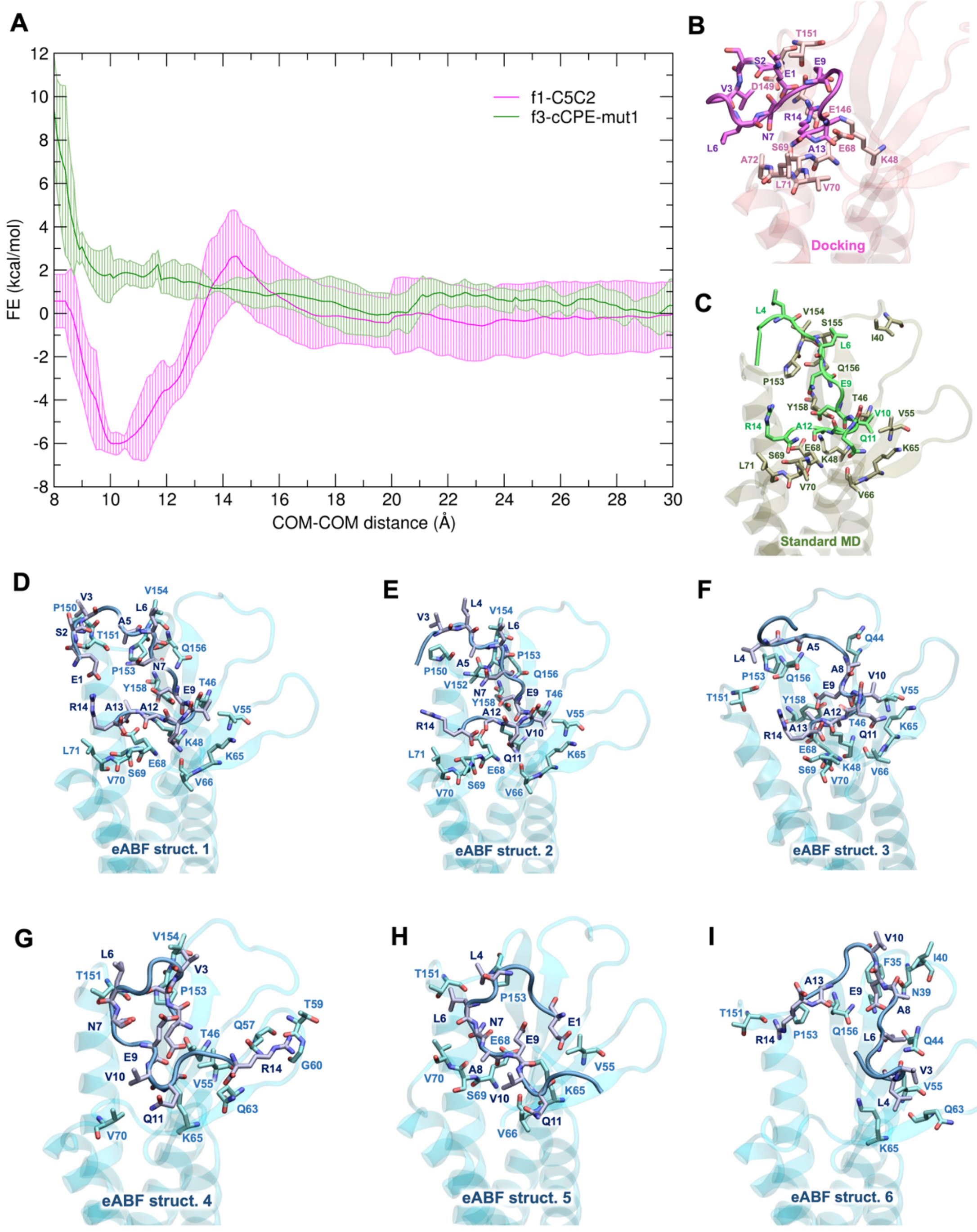


***Figure S7. Unbinding free energy calculation for the CLDN5-peptide complexes calculated with eABF.*** ***A.*** *Free energy (FE) profiles for the f1-C5C2 (magenta) and f3-cCPE-mut1 (green) are computed with respect to the distance between the peptides and the CLDN5 ECL COMs. FE estimate and errors are indicated as the mean ± SD for three 150 ns-long replicates. Representative structures of the mCLDN5-peptide complex are shown for window1 (spanning from 8 to 20 Å) and window2 (spanning from 20 to 30 Å) and colored in magenta and green for f1-C5C2 and f3-cCPE-mut1, respectively.* ***B-I.*** *Individual binding configurations provided by docking (**B**), standard MD (**C**) and representative structures resulting from clustering of eABF trajectory (**D-I**). Amino acids involved in the mCLDN5-f1-C5C2 interactions are shown as sticks.*

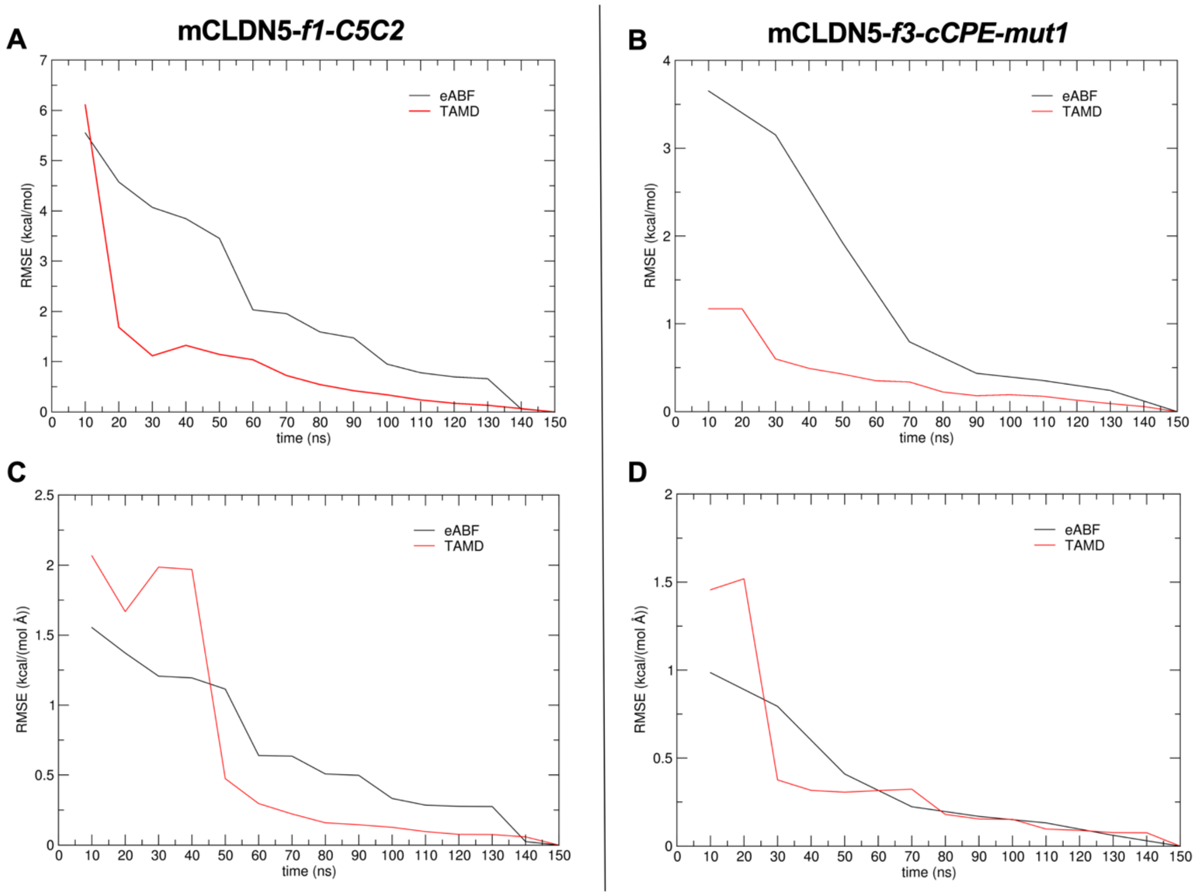


***Figure S8. Convergence analysis of eABF and TAMD calculations.*** *The convergence was assessed by computing the root-mean square error (RMSE) of the estimated FE profile (**A**, **B**) or the FE gradients (**C**, **D**) every 10 ns of simulation with respect to the FE profile or FE gradients at 150 ns. The analysis was conducted for a representative replicate of the mCLDN5-f1-C5C2 (**A**, **C**) and mCLDN5-f3-cCPE-mut1 (**B**, **D**) systems simulated with the TAMD (red line) and eABF (black line) algorithms.*

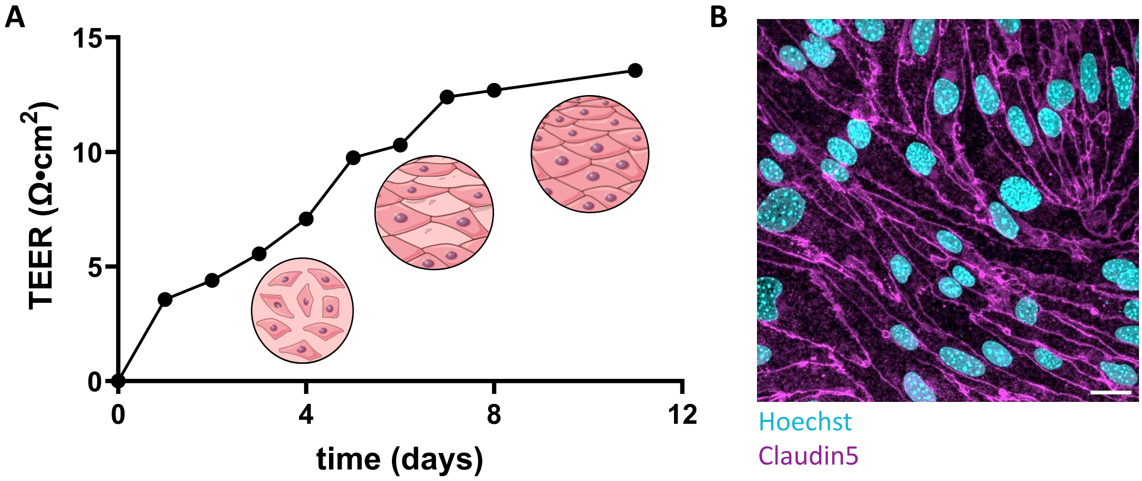


***Figure S9. Characterization of the 2D BBB model based on bEND.3 cells cultured on Transwell membranes (0.4 µm pore size)*. *A.* The recordings of TEER values as a function of the days in vitro (DIV) show a progressive increase due to the formation of the tight endothelial cell barrier. *B.* Representative confocal image of CLDN5 immunostaining at TJs. Scale bar, 20 µm.***

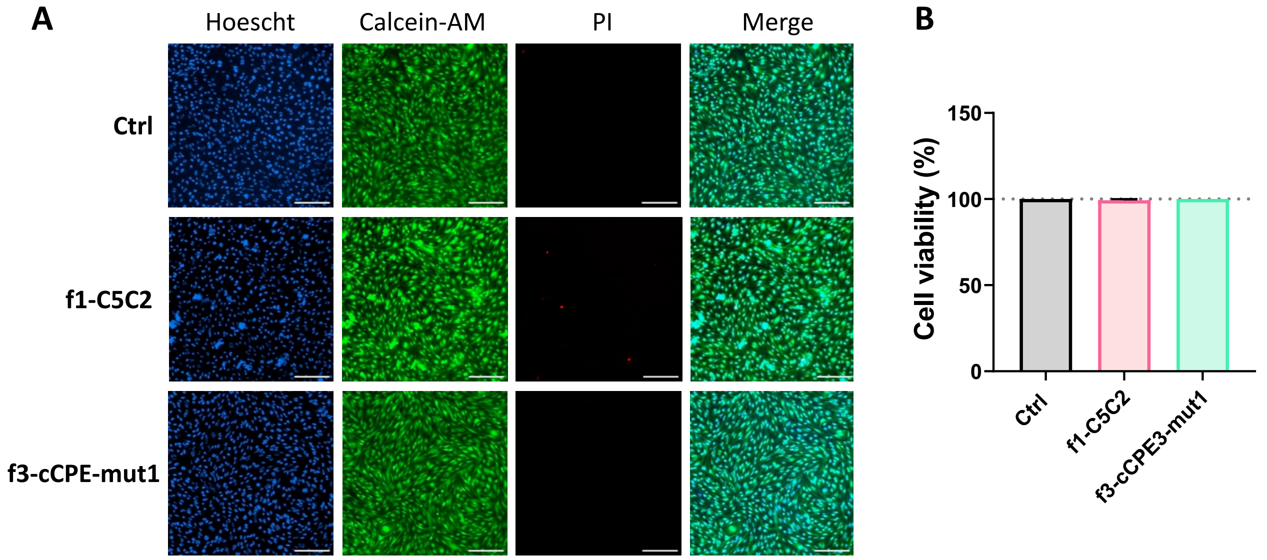


***Figure S10*. Viability test for bEnd.3 cells exposed to f3-cCPE-mut1 and f1-C5C2 (25 µM) for 24 h. A.** *Representative fluorescence images for cells stained with Hoechst (nuclear staining), Calcein-AM (viable cells), and Propidium iodide (PI – dead cells).* ***B****. Quantification of the mean (± sem) percentage of viable cells over total cells (% cell viability) for bEnd.3 cells treated with either vehicle (Ctrl) or the selected peptides. Scale bars, 100 µm.*

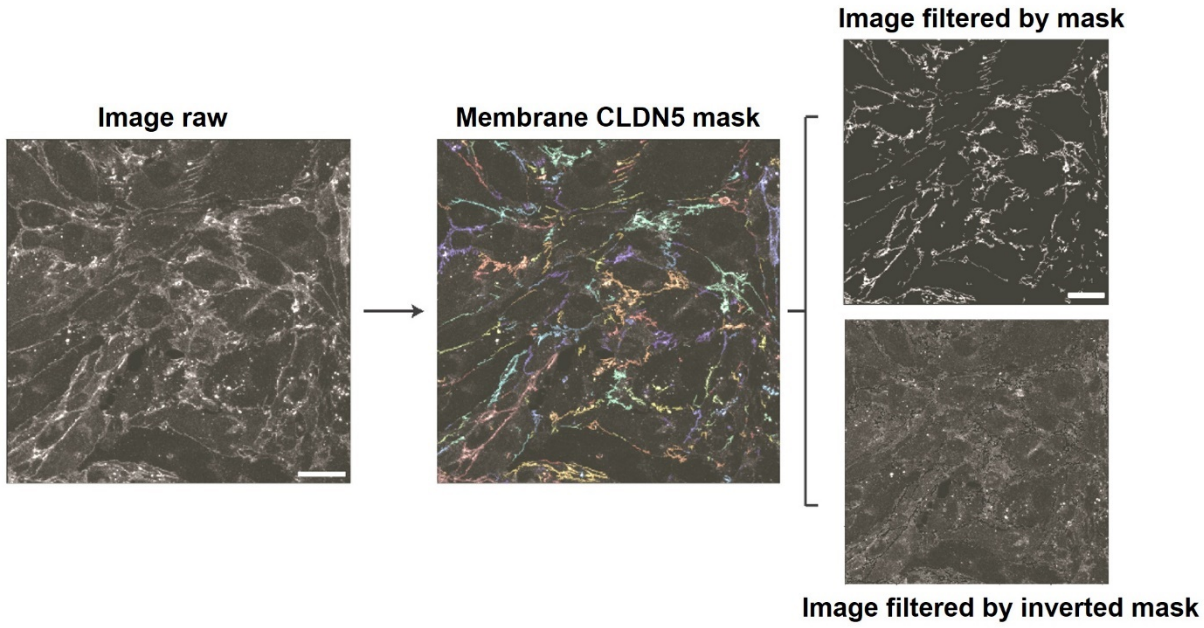


***Figure S11. Schematic workflow of CLDN5 image analysis on CellProfiler.*** *A membrane CLDN5 mask was built starting from raw maximal projection images. Each image was then filtered by both CLDN5 mask and inverted CLDN5 mask to obtain and quantify only membrane or cytosolic fluorescence intensity. Scale bars, 25 μm.*

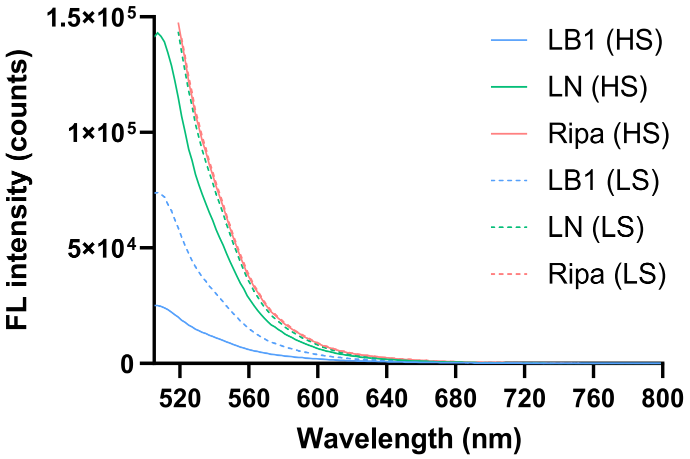


***Figure S12. Optimization of the cell lysis method.*** *Three lysis protocols were compared: (1) Lysis Buffer 1 (LB1 - 20 mM Tris/HCl, pH 7.5, 100 mM NaCl, 1 mM $MgCl_2$, 5% glycerol and protease inhibitor cocktail), (2) Lysis Buffer 2 (RIPA – 50 mM Tris HCL pH 7.4, 50 mM NaCl, 2 mM EDTA, 0.1% SDS and protease inhibitor cocktail), (3) flash freeze-thawing in liquid nitrogen (LN). The protocol was ended with centrifugation at either low (LS; 2,800 x g) or high (HS; 16,200 x g) speed. The GFP emission spectra of the various lysates are reported.*

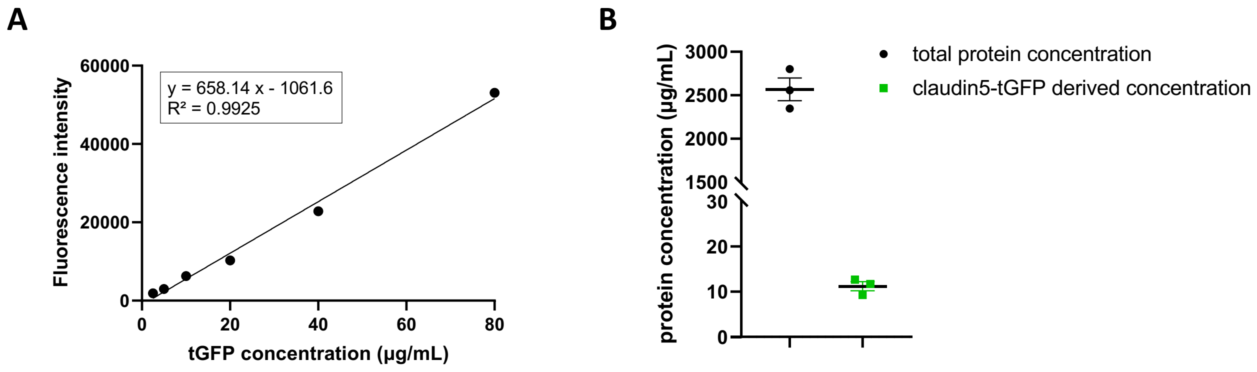


***Figure S13. Quantification of mCLDN5-tGFP from HEK293T cell lysates. A.*** *A calibration curve was derived by diluting increasing concentrations of recombinant tGFP in PBS and measuring the resulting fluorescence intensity.* ***B.*** *The mCLDN5-tGFP concentration from the various cell lysates was derived and compared to the total amount of protein obtained by direct quantitation using the Nanodrop A280 method.*

### Caption to the Supplementary Video

***Video S1. Trajectories of the binding of f1-C5C2 and f3-cCPE-mut1 to mCLDN5.*** *In the video, mCLDN5 is colored in pink and it is embedded in the rectangular lipid membrane shown in transparency. Peptides backbones are represented as cyan and green sticks for f1-C5C2 and f3-cCPE-mut1, respectively, whereas N atoms are colored in blue and O atoms in red. Water and ions are not shown for clarity.*

### Supplementary Tables

##### *Table S1. List of the peptides studied in this work*

| PEPTIDE | SEQUENCE | GRAVY (hydrophilic < 0) | Residues numbering in the entire protein | Predicted solubility according to KNN model | Solubility descriptors | | | Ref.s |
|---|---|---|---|---|---|---|---|---|
| | | | | | Largest aggregate | Number of aggregates | Water molecules contacts at the end of the simulation (%) | |
| **f1-cCPE** | **AGNLYDWRSSN** | -1.236 | 220 - 230 | Insoluble | 20.01 | 4.79 | 58.11 | [2,67] |
| f1-cCPE-mut1 | RGNLYDWRSDN | -2.055 | | Insoluble | 13.00 | 3.88 | 81.85 | |
| f1-cCPE-mut2 | GGNLYDYRSSN | -1.473 | | Insoluble | 11.65 | 3.50 | 73.19 | |
| **f2-cCPE** | **SWSGNYPYHILFQKF** | -0.487 | 305 - 319 | Insoluble | 20.71 | 2.81 | 65.05 | [2,67] |
| f2-cCPE-mut1 | SWSDNYQYDTRFQKF | -1.740 | | Insoluble | 14.94 | 4.00 | 76.98 | |
| **f3-cCPE** | **ATERLNLTDALNSNPA** | -0.562 | 205 - 220 | Soluble | 6.38 | 8.94 | 82.82 | [2,67] |
| f3-cCPE-mut1 | RTERLDKTDASNSNPA | -1.725 | | Soluble | 9.84 | 11.33 | 76.02 | |
| **f1-C5C2** | **ESVLALNAEVQAAR** | 0.279 | 68 - 81 | Soluble | 7.19 | 11.05 | 75.51 | [6,7] |
| **f2-C5C2** | **VLALNAEVQAAR** | 0.683 | 70 - 81 | Insoluble | 11.44 | 4.26 | 64.76 | |
| f1-C5C2-mut1 | DSVLALSTEVQKSR | -0.300 | | Insoluble | 15.66 | 11.38 | 82.82 | |
| f1-C5C2-mut2 | DSVLALSTQVQAAR | 0.293 | | Insoluble | 14.12 | 7.48 | 91.71 | |
| f1-C5C2-mut3 | ESVYALNAKVQDAR | -0.493 | | Soluble | 8.00 | 9.12 | 94.16 | |
| f1-C5C2-mut4 | ESVLATNAQVQSAR | -0.229 | | Insoluble | 9.04 | 7.37 | 82.06 | |
| f1-C5C2-mut5 | ESVLALNAEVQKSR | -0.314 | | Insoluble | 14.00 | 6.63 | 87.80 | |
| f1-C5C2-mut6 | ESVLALSAEVQAAR | -0.314 | | Soluble | 5.46 | 10.02 | 91.78 | |
| f1-C5C2-mut7 | DSVLALSTEVQAAR | 0.293 | | Insoluble | 18.42 | 6.12 | 76.05 | |
| f1-C5C2-mut8 | DSVLALNTEVQAAR | 0.100 | | Soluble | 11.82 | 10.18 | 78.27 | |

*The already published fragments of either cCPE or CLDN5 ECL1 are indicated in the first row of each block. Mutations introduced in each sequence are indicated with the color-code based on their charge: blue for basic, red for acidic and green for neutral amino acids, and the GRAVY coefficient of each peptide is indicated. The size of the largest aggregate and the number of aggregates is reported as average values over the 10 ns of MD simulation. The percentage of water molecules contacts at the end of the simulation is calculated with respect to the initial configuration. The peptides studied in this work are highlighted in orange.*

#### *Table S2. Peptides external dataset for the solubility classification*

| Name | Code NAME | SEQUENCE | GRAVY (hydrophilic < 0) | largest aggregate | number of aggregates | Water molecules contacts at the end of the simulation (%) |
|---|---|---|---|---|---|---|
| **Temporin-A** | Pep_Ins1 | FLPLIGRVLSGIL | 1.808 | 12.00 | 5.00 | 60.15 |
| **Temporin-B** | Pep_Ins2 | LLPIVGNLLKSLL | 1.638 | 10.00 | 4.00 | 51.22 |
| **Crabrolin** | Pep_Ins3 | FLPLILRKIVTAL | 1.715 | 9.00 | 5.00 | 60.12 |
| **Uperin 7.1** | Pep_Ins4 | GWFDVVKHIASAV | 0.831 | 12.00 | 4.00 | 54.73 |
| **Polybia-MP-III** | Pep_Ins5 | IDWLKLGKMVMDVL | 0.860 | 15.00 | 8.00 | 60.33 |
| **Peptide A1** | Pep_Ins6 | FLPAIAGILSQLF | 1.792 | 21.00 | 2.00 | 40.92 |
| **Meucin-13** | Pep_Ins7 | IFGAIAGLLKNIF | 1.700 | 14.00 | 5.00 | 58.61 |
| **Ranatuerin 9** | Pep_Ins8 | FLFPLITSFLSKVL | 1.750 | 15.00 | 2.00 | 56.40 |
| **Alloferon-2** | Pep_Sol1 | GVSGHGQHGVHG | -0.625 | 7.00 | 17.00 | 89.15 |
| **Histone H6-like protein** | Pep_Sol2 | PKRKSATKGDEPA | -1.900 | 5.49 | 15.58 | 86.75 |
| **Crinicepsin-2** | Pep_Sol3 | RERSKGSKYLYVG | -1.331 | 6.00 | 14.00 | 76.66 |
| **A21978C1** | Pep_Sol4 | WNDTGKDADGSEY | -1.854 | 6.74 | 15.34 | 83.53 |
| **human Histatine-8** | Pep_Sol5 | KFHEKHHSHRGY | -2.358 | 7.50 | 14.00 | 71.21 |
| **Astracidin-2** | Pep_Sol6 | RPRPNYRPRPIYRD | -2.429 | 4.00 | 13.00 | 77.70 |
| **HRNR1132-1143** | Pep_Sol7 | GSGSRQSPSYGR | -1.650 | 4.00 | 16.00 | 84.78 |
| **Odorranain-R1** | Pep_Sol8 | GFSPNLPGKGLRIS | -0.214 | 7.50 | 11.50 | 70.35 |

*Soluble (8) and insoluble (8) peptides were selected from the* ***aps.unmc.edu*** *database. MD simulations for the solubility assessment were performed with the same protocol described for the mCLDN5-binding peptides. The following parameters were computed: dimension of the largest aggregate, number of aggregates and percentage of water molecule contacts between the peptides and the solvent at the end of the simulation with respect to the starting configuration.*

***Table S3. Analysis of the interactions identified in mCLDN5-f1-C5C2 binding configuration resulting from molecular docking, standard MD and biased MD (TAMD, eABF) simulations.***

| *Interaction (mCLDN5 – f1-C5C2)* | *Type of interaction* | *Bond length (Å)* | | | | | | | |
|---|---|---|---|---|---|---|---|---|---|
| | | **Docking** | **Standard MD** | **TAMD struct. 1 (RMSD: 2.32 Å)** | **TAMD struct. 2 (RMSD: 3.32 Å)** | **TAMD struct. 3 (RMSD: 5.93 Å)** | **TAMD struct. 4 (RMSD: 9.24 Å)** | **TAMD struct. 5 (RMSD: 12.07 Å)** | **TAMD struct. 6 (RMSD: 14.50 Å)** |
| | | | | **eABF struct. 1 (RMSD: 1.94 Å)** | **eABF struct. 2 (RMSD: 2.91 Å)** | **eABF struct. 3 (RMSD: 5.14 Å)** | **eABF struct. 4 (RMSD: 9.54 Å)** | **eABF struct. 5 (RMSD: 11.01 Å)** | **eABF struct. 6 (RMSD: 14.56 Å)** |
| T33(OG1) – E9(OE1) | HB | | | | | | | 2.68 | 2.69 |
| F35 (CD2) – E9 (CG) | apolar | | | | | | | | 3.42 |
| I40 (CG1) – L6 (CD1) | apolar | | | | | 3.86 | | | |
| T42 (O) – N7 (ND2) | HB | | | | | | 2.95 | | |
| Q44 (NE2) – L6 (O) | HB | | | | | | | | 2.98 |
| Q44(CA) – A8(CB) | apolar | | | | | | | 3.92 | |
| Q44(OE1) – A8(O) | HB | | | | | | | | 3.43 |
| T46 (OG1) – A8 (N) | HB | | | | | | | 3.88 | |
| T46 (CG2) – E9 (CB) | apolar | | | 3.96 | | | | | |
| T46 (CG2) – V10 (CG2) | apolar | | | | 3.50 | | | | |
| K48 (NZ) – R14 (NH2) | HB | 3.46 | | | | | | | |
| K48 (NZ) – E9 (OE2) | SB | | 2.63 | 2.64 | 2.68 | 2.61 | | 2.67 | 2.61 |
| V55 (CG1) – L6 (CA) | apolar | | | | | | | | 3.54 |
| V55 (CG1) – N7 (CA) | apolar | | | | | | | 3.92 | |
| V55 (CG1) – V10 (CG2) | apolar | | | 3.55 | 3.84 | 3.72 | | | |
| V55 (CG1) – V10 (CB) | apolar | | | | 3.91 | 3.94 | | | |
| Q57 (OE1) – A5 (N) | HB | | | | | | | | 2.88 |
| Q57 (NE2) – A5 (O) | HB | | | | | | | 3.02 | |
| Q57 (NE2) – R14 (NE) | HB | | | | | | 3.41 | | |
| S58(OG) – V8(O) | HB | | | | | | | | 3.65 |
| T59(CG2) – V3(CG2) | apolar | | | | | | | | 3.93 |
| Q63 (CB) – L4 (CD2) | apolar | | | | | | | | 3.74 |
| Q63 (NE2) – R14 (OT2) | HB | | | | | | 3.35 | | |
| K65 (NZ) – E1 (OE1) | SB | | | | | | 2.66 | | |
| K65 (NZ) – L6 (O) | HB | | | | | | | | 2.73 |
| K65 (NZ) – E9 (O) | HB | | | | | | 2.83 | | |
| K65 (CG) – V10 (CG2) | apolar | | | | | 3.77 | | | |
| V66 (O) – Q11 (NE2) | HB | | | 2.72 | 2.83 | | | | |
| Y67 (O) – Q11(OE1) | HB | | | | 3.70 | | | | |
| E68 (OE2) – N7 (ND2) | HB | | | | | | | 3.75 | |
| E68 (N) – Q11 (OE1) | HB | | | 3.65 | 3.48 | | | | |
| E68 (CD) – A13 (CB) | apolar | 2.87 | | | | | | | |
| E68 (OE2) – A13 (O) | VdW | 3.21 | | 3.46 | | | | | |
| E68 (CB) – A12 (CB) | apolar | | 3.12 | 3.85 | 3.78 | 3.90 | | | |
| E68(OE1) – A12 (N) | HB | | | | | 3.97 | | | |
| E68(OE1) – A12 (O) | VdW | | | | | 3.99 | | | |
| S69 (N) – E1 (O) | HB | | | | | | 3.46 | | |
| S69 (OG) – N7 (OD1) | HB | 3.45 | | | | | | | |
| S69 (N) – A8 (O) | HB | | | | | | | 3.97 | |
| S69 (O) – A8 (N) | HB | | | | | | | 2.94 | |
| S69 (N) – A12 (O) | HB | | 3.04 | 3.17 | 2.81 | 3.50 | | | |
| S69 (N) – A13 (O) | HB | | | | 3.91 | 3.37 | | | |
| S69 (OG) – A13 (O) | HB | | | | | 3.88 | | | |
| S69 (CB) – A13 (C) | apolar | | 4.00 | | | | | | |
| S69 (CA) – A13 (C) | apolar | | | | | 3.83 | | | |
| S69 (N) – R14 (N) | HB | 2.99 | | | | | | | |
| S69 (OG) – R14 (N) | HB | | 3.29 | | 3.05 | 3.19 | | | |
| S69 (OG) – R14 (O) | HB | | 2.84 | | | | | | |
| V70 (N) – R14 (N) | HB | 3.74 | | | | | | | |
| V70 (N) – R14 (O) | HB | | 2.74 | 2.84 | 2.93 | 2.92 | 3.02 | | |
| V70 (CG1) – R14 (CB) | apolar | | | | 3.98 | | | | |
| L71 (CG) – R14 (CB) | apolar | | | | 3.81 | 3.94 | | | |
| L71 (N) – R14 (O) | HB | | 3.15 | | | | | | |
| P150 (O) – E1 (N) | HB | | | | | 2.55 | | | |
| P150 (O) – L4 (N) | HB | | | | 3.01 | | | | |
| T151 (CG2) – E1 (CB) | apolar | | | 3.40 | | | | | |
| T151 (OG1) - E1 (OE1) | HB | 2.86 | | | | | | | |
| T151 (O) – L4 (N) | HB | | | | 3.52 | | | | |
| T151 (OG1) – E9 (OE1) | HB | 3.22 | | | | | | | |

| | | | | | | | | |
|---|---|---|---|---|---|---|---|---|
| T151 (O) N7 (ND2) | HB | | | | | 2.89 | | |
| T151 (O) – R14 (NH2) | HB | | | | | | | 2.77 |
| V152 (C) – E1 (CG) | apolar | | | | 3.66 | | | |
| V152 (O) – A5 (N) | HB | | | 3.80 | | | | |
| P153 (CD)- L4 (CD2) | apolar | | | | | | 3.87 | |
| P153 (CG) – A5 (CB) | apolar | | 3.87 | | | | | |
| P153 (CB) – A5 (C) | apolar | | | | | 3.94 | | |
| P153 (CB) – N7 (CB) | apolar | 3.66 | 3.87 | | | | | |
| P153 (CG) – N7 (CG) | apolar | 3.76 | 3.95 | | | | | |
| P153 (CB) – V10 (CG2) | apolar | | | | | | 3.51 | |
| V154 (CG2) – V3 (CG1) | apolar | | | | | 3.85 | | |
| V154 (N) – A5 (O) | HB | 3.25 | 3.12 | 3.16 | | | | |
| V154 (CG2) – L4 (C) | apolar | 3.93 | | | | | | |
| V154 (CG1) – L6 (CD2) | apolar | | 3.60 | | | | | |
| Q156 (OE1) – N7 (ND2) | HB | | 2.91 | | | | | |
| Q156 (NE2) – A8 (O) | HB | | | | | | 2.67 | 2.96 |
| Q156 (NE2) – E9 (OE1) | HB | | | 3.48 | | | | 3.46 |
| Y158 (OH) – E9 (OE2) | HB | 2.72 | 2.89 | 3.76 | 2.64 | | | |

*Representative structures from TAMD and eABF trajectories were selected from a RMSD-based clustering over the configurations characterized by a CV value consistent with the energy minimum. RMSD was calculated with respect to the peptide pose resulting from standard MD simulation. Atoms involved in each interaction are labelled according to the CHARMM atom type. A legend for the atom types and the corresponding atom in the amino acids is provided in **Table S4**. Interactions observed in at least two structures are highlighted.*

***Table S4. Definition of the CHARMM atom types for the interactions included in Table S3.***

| CHARMM atom type | Corresponding atom in the amino acid |
|---|---|
| C | backbone carbonyl carbon |
| CG | carbon gamma |
| CG1 | secondary carbon gamma in isoleucine or primary carbon gamma in valine |
| CG2 | primary carbon gamma in isoleucine and valine |
| CB | carbon beta |
| CD | carbon delta |
| CD1/2 | carbon delta in leucine |
| N | backbone amidic nitrogen |
| NE | nitrogen in secondary amine in guanidine group of arginine |
| NE2 | amidic nitrogen in glutamine |
| NZ | terminal charged nitrogen in lysine |
| NH2 | nitrogen in primary neutral amine in guanidine group of arginine |
| ND2 | amidic nitrogen in asparagine |
| O | backbone carbonyl oxygen |
| OE1/2 | carboxyl oxygen in glutamate and glutamine (OE1) |
| OG | hydroxyl oxygen in serine |
| OG1 | hydroxyl oxygen in threonine |
| OD1/2 | carboxyl oxygen in aspartate and asparagine (OD1) |
| OH | aromatic hydroxyl oxygen |
| OT1/2 | terminal carboxyl oxygen |

### Supplementary references


1. Wang G, Li X, Wang Z (2016) APD3: the antimicrobial peptide database as a tool for research and education. *Nucleic Acids Res* 44: D1087-1093.
2. Neuhaus W, Piontek A, Protze J, et al. (2018) Reversible opening of the blood-brain barrier by claudin-5-binding variants of Clostridium perfringens enterotoxin's claudin-binding domain. *Biomaterials* 161: 129–143.
3. Shinoda T, Shinya N, Ito K, et al. (2016) Structural basis for disruption of claudin assembly in tight junctions by an enterotoxin. *Sci Rep* 6: 33632.
4. Zhang J, Liang Y, Zhang Y (2011) Atomic-Level Protein Structure Refinement Using Fragment-Guided Molecular Dynamics Conformation Sampling. *Structure* 19: 1784–1795.
5. Gasteiger E, Hoogland C, Gattiker A, et al. (2005) Protein Identification and Analysis Tools on the ExPASy Server, In: Walker JM (Ed.), *The Proteomics Protocols Handbook*, Totowa, NJ, Humana Press, 571–607.
6. Dithmer S, Staat C, Müller C, et al. (2017) Claudin peptidomimetics modulate tissue barriers for enhanced drug delivery. *Ann N Y Acad Sci* 1397: 169–184.
7. Staat C, Coisne C, Dabrowski S, et al. (2015) Mode of action of claudin peptidomimetics in the transient opening of cellular tight junction barriers. *Biomaterials* 54: 9–20.
8. Waterhouse A, Bertoni M, Bienert S, et al. (2018) SWISS-MODEL: homology modelling of protein structures and complexes. *Nucleic Acids Res* 46: W296–W303.
9. Suzuki H, Nishizawa T, Tani K, et al. (2014) Crystal structure of a claudin provides insight into the architecture of tight junctions. *Science* 344: 304–307.
10. Alberini G, Benfenati F, Maragliano L (2017) A refined model of claudin-15 tight junction paracellular architecture by molecular dynamics simulations. *PLOS ONE* 12: e0184190.
11. Suzuki H, Tani K, Tamura A, et al. (2015) Model for the Architecture of Claudin-Based Paracellular Ion Channels through Tight Junctions. *J Mol Biol* 427: 291–297.
12. Irudayanathan FJ, Wang N, Wang X, et al. (2017) Architecture of the paracellular channels formed by claudins of the blood-brain barrier tight junctions. *Ann N Y Acad Sci* 1405: 131–146.
13. Piontek J, Krug SM, Protze J, et al. (2020) Molecular architecture and assembly of the tight junction backbone. *Biochim Biophys Acta BBA - Biomembr* 1862: 183279.
14. Berselli A, Alberini G, Benfenati F, et al. (2022) Computational Assessment of Different Structural Models for Claudin-5 Complexes in Blood–Brain Barrier Tight Junctions. *ACS Chem Neurosci* 13: 2140–2153.
15. Berselli A, Alberini G, Benfenati F, et al. (2022) Computational study of ion permeation through claudin-4 paracellular channels. *Ann N Y Acad Sci* 1516: 162–174.
16. Hurwitz N, Schneidman-Duhovny D, Wolfson HJ (2016) Memdock: an α-helical membrane protein docking algorithm. *Bioinforma Oxf Engl* 32: 2444–2450.
17. Humphrey W, Dalke A, Schulten K (1996) VMD: visual molecular dynamics. *J Mol Graph* 14: 33–38, 27–28.
18. Heo L, Lee H, Seok C (2016) GalaxyRefineComplex: Refinement of protein-protein complex model structures driven by interface repacking. *Sci Rep* 6: 32153.
19. Lamiable A, Thévenet P, Rey J, et al. (2016) PEP-FOLD3: faster de novo structure prediction for linear peptides in solution and in complex. *Nucleic Acids Res* 44: W449–W454.
20. Still WC, Tempczyk A, Hawley RC, et al. (1990) Semianalytical treatment of solvation for molecular mechanics and dynamics. *J Am Chem Soc* 112: 6127–6129.
21. Constanciel R, Contreras R (1984) Self consistent field theory of solvent effects representation by continuum models: Introduction of desolvation contribution. *Theor Chim Acta* 65: 1–11.
22. Jorgensen WL, Chandrasekhar J, Madura JD, et al. (1983) Comparison of simple potential functions for simulating liquid water. *J Chem Phys* 79: 926–935.
23. Jo S, Kim T, Iyer VG, et al. (2008) CHARMM-GUI: a web-based graphical user interface for CHARMM. *J Comput Chem* 29: 1859–1865.
24. Feller SE, Zhang Y, Pastor RW, et al. (1995) Constant pressure molecular dynamics simulation: The Langevin piston method. *J Chem Phys* 103: 4613–4621.
25. Martyna GJ, Tobias DJ, Klein ML (1994) Constant pressure molecular dynamics algorithms. *J Chem Phys* 101: 4177–4189.
26. Darden T, York D, Pedersen L (1993) Particle mesh Ewald: An N·log(N) method for Ewald sums in

large systems. *J Chem Phys* 98: 10089–10092.
27. Steinbach PJ, Brooks BR (1994) New spherical-cutoff methods for long-range forces in macromolecular simulation. *J Comput Chem* 15: 667–683.
28. Ryckaert J-P, Ciccotti G, Berendsen HJC (1977) Numerical integration of the cartesian equations of motion of a system with constraints: molecular dynamics of n-alkanes. *J Comput Phys* 23: 327–341.
29. Miyamoto S, Kollman PA (1992) SETTLE: an analytical version of the SHAKE and RATTLE algorithm for rigid water models. *J Comput Chem* 13: 952–962.
30. Phillips JC, Hardy DJ, Maia JDC, et al. (2020) Scalable molecular dynamics on CPU and GPU architectures with NAMD. *J Chem Phys* 153: 044130.
31. Huang J, Rauscher S, Nawrocki G, et al. (2017) CHARMM36m: an improved force field for folded and intrinsically disordered proteins. *Nat Methods* 14: 71–73.
32. Kuroda Y, Suenaga A, Sato Y, et al. (2016) All-atom molecular dynamics analysis of multi-peptide systems reproduces peptide solubility in line with experimental observations. *Sci Rep* 6: 19479.
33. Xu D, Zhang Y (2011) Improving the physical realism and structural accuracy of protein models by a two-step atomic-level energy minimization. *Biophys J* 101: 2525–2534.
34. Jo S, Cheng X, Islam SM, et al. (2014) CHARMM-GUI PDB manipulator for advanced modeling and simulations of proteins containing nonstandard residues. *Adv Protein Chem Struct Biol* 96: 235–265.
35. Klauda JB, Venable RM, Freites JA, et al. (2010) Update of the CHARMM all-atom additive force field for lipids: validation on six lipid types. *J Phys Chem B* 114: 7830–7843.
36. Trott O, Olson AJ (2010) AutoDock Vina: Improving the speed and accuracy of docking with a new scoring function, efficient optimization, and multithreading. *J Comput Chem* 31: 455–461.
37. Goodsell DS, Sanner MF, Olson AJ, et al. (2021) The AutoDock suite at 30. *Protein Sci Publ Protein Soc* 30: 31–43.
38. Morris GM, Huey R, Lindstrom W, et al. (2009) AutoDock4 and AutoDockTools4: Automated docking with selective receptor flexibility. *J Comput Chem* 30: 2785–2791.
39. Pettersen EF, Goddard TD, Huang CC, et al. (2004) UCSF Chimera--a visualization system for exploratory research and analysis. *J Comput Chem* 25: 1605–1612.
40. London N, Raveh B, Cohen E, et al. (2011) Rosetta FlexPepDock web server--high resolution modeling of peptide-protein interactions. *Nucleic Acids Res* 39: W249-253.
41. Alam N, Schueler-Furman O (2017) Modeling Peptide-Protein Structure and Binding Using Monte Carlo Sampling Approaches: Rosetta FlexPepDock and FlexPepBind. *Methods Mol Biol Clifton NJ* 1561: 139–169.
42. Ciemny M, Kurcinski M, Kamel K, et al. (2018) Protein–peptide docking: opportunities and challenges. *Drug Discov Today* 23: 1530–1537.
43. Rentzsch R, Renard BY (2015) Docking small peptides remains a great challenge: an assessment using AutoDock Vina. *Brief Bioinform* 16: 1045–1056.
44. Noskov SY, Roux B (2008) Control of Ion Selectivity in LeuT: Two Na+ Binding Sites with two different mechanisms. *J Mol Biol* 377: 804–818.
45. Luo Y, Roux B (2010) Simulation of Osmotic Pressure in Concentrated Aqueous Salt Solutions. *J Phys Chem Lett* 1: 183–189.
46. Venable RM, Luo Y, Gawrisch K, et al. (2013) Simulations of anionic lipid membranes: development of interaction-specific ion parameters and validation using NMR data. *J Phys Chem B* 117: 10183–10192.
47. Fiorin G, Klein ML, Hénin J (2013) Using collective variables to drive molecular dynamics simulations. *Mol Phys* 111: 3345–3362.
48. Maragliano L, Vanden-Eijnden E (2006) A temperature accelerated method for sampling free energy and determining reaction pathways in rare events simulations. *Chem Phys Lett* 426: 168–175.
49. Stoltz G, Vanden-Eijnden E (2018) Longtime convergence of the temperature-accelerated molecular dynamics method. *Nonlinearity* 31: 3748.
50. Lesage A, Lelièvre T, Stoltz G, et al. (2017) Smoothed Biasing Forces Yield Unbiased Free Energies with the Extended-System Adaptive Biasing Force Method. *J Phys Chem B* 121: 3676–3685.
51. Fu H, Shao X, Chipot C, et al. (2016) Extended Adaptive Biasing Force Algorithm. An On-the-Fly Implementation for Accurate Free-Energy Calculations. *J Chem Theory Comput* 12: 3506–3513.
52. Comer J, Gumbart JC, Hénin J, et al. (2015) The Adaptive Biasing Force Method: Everything You Always Wanted To Know but Were Afraid To Ask. *J Phys Chem B* 119: 1129–1151.
53. Woo H-J, Roux B (2005) Calculation of absolute protein–ligand binding free energy from computer simulations. *Proc Natl Acad Sci U S A* 102: 6825–6830.

54. Blazhynska M, Goulard Coderc de Lacam E, Chen H, et al. (2022) Hazardous Shortcuts in Standard Binding Free Energy Calculations. *J Phys Chem Lett* 13: 6250–6258.
55. Allen TW, Andersen OS, Roux B (2004) Energetics of ion conduction through the gramicidin channel. *Proc Natl Acad Sci U S A* 101: 117–122.
56. Roux B, Andersen OS, Allen TW (2008) Comment on "Free energy simulations of single and double ion occupancy in gramicidin A" [J. Chem. Phys. 126, 105103 (2007)]. *J Chem Phys* 128: 227101.
57. Gumbart JC, Roux B, Chipot C (2013) Standard Binding Free Energies from Computer Simulations: What Is the Best Strategy? *J Chem Theory Comput* 9: 794–802.
58. Gumbart JC, Roux B, Chipot C (2013) Efficient Determination of Protein–Protein Standard Binding Free Energies from First Principles. *J Chem Theory Comput* 9: 3789–3798.
59. Fu H, Cai W, Hénin J, et al. (2017) New Coarse Variables for the Accurate Determination of Standard Binding Free Energies. *J Chem Theory Comput* 13: 5173–5178.
60. Limongelli V, Bonomi M, Parrinello M (2013) Funnel metadynamics as accurate binding free-energy method. *Proc Natl Acad Sci U S A* 110: 6358–6363.
61. Limongelli V (2020) Ligand binding free energy and kinetics calculation in 2020. *WIREs Comput Mol Sci* 10.
62. Fu H, Chen H, Blazhynska M, et al. (2022) Accurate determination of protein:ligand standard binding free energies from molecular dynamics simulations. *Nat Protoc* 17: 1114–1141.
63. Lan J, Ge J, Yu J, et al. (2020) Structure of the SARS-CoV-2 spike receptor-binding domain bound to the ACE2 receptor. *Nature* 581: 215–220.
64. Lapelosa M (2018) Conformational dynamics and free energy of BHRF1 binding to Bim BH3. *Biophys Chem* 232: 22–28.
65. Lapelosa M (2017) Free Energy of Binding and Mechanism of Interaction for the MEEVD-TPR2A Peptide–Protein Complex. *J Chem Theory Comput* 13: 4514–4523.
66. Doudou S, Burton NA, Henchman RH (2009) Standard Free Energy of Binding from a One-Dimensional Potential of Mean Force. *J Chem Theory Comput* 5: 909–918.
67. Liao Z, Yang Z, Piontek A, et al. (2016) Specific binding of a mutated fragment of Clostridium perfringens enterotoxin to endothelial claudin-5 and its modulation of cerebral vascular permeability. *Neuroscience* 327: 53–63.
